# Defining the SPCA2C interactome identifies unique links to Store-Operated Ca^2+^ Entry

**DOI:** 10.1101/2023.09.13.557610

**Authors:** Petra Samardzija, Melissa A Fenech, Ryann Lang, McKenzie C Carter, Stephanie Chen, Selina Shi, Peter B Stathopulos, Christopher L Pin

## Abstract

Calcium (Ca^2+^) is critical for normal cell function and several protein networks are required for Ca^2+^ signaling. In the pancreas, regulated changes in cytosolic Ca^2+^ allow for the exocytosis of zymogen granules and altered Ca^2+^ signaling underlies pancreatic pathologies. Previously, our laboratory showed a pancreas-specific isoform of secretory pathway Ca^2+^-ATPase 2 (SPCA2C) affects multiple pathways involved in Ca^2+^ homeostasis. The goal of this study was to define the SPCA2C interactome that contributes to these processes. Using proximity-dependent biotin identification, BioID, we expressed SPCA2C-BirA^*HA^ in HEK293 cells with constitutive Orai1 expression. *In silico* modeling of SPCA2C showed a highly dynamic cytosolic C-terminus, revealing a putative site for interactions and was selected for BirA* fusion. 150 candidate SPCA2C interactors were identified. Gene Ontology and KEGG Pathway analyses supported localization of SPCA2C to the endoplasmic reticulum as well as function in Ca^2+^ signaling and suggested roles in vesicular transport. SPCA2C interactions with stromal interaction molecule 1 (STIM1), extended synaptotagmin-1 (ESYT1), and aspartate beta-hydroxylase (ASPH) were validated. Coiled-coil domain containing protein 47 (CCDC47) ranked as a high confidence SPCA2C-interactor which was confirmed through immunoprecipitation and co-localization with SPCA2C. CCDC47 interactions with SPCA2C, STIM1 and Orai1 were maintained following deletion of the CCDC47 luminal and cytosolic domains while deletion of only the coiled-coil domain altered CCDC47 localization and decreased interactions with STIM1 and Orai1. Overall, this study defines several novel protein interactions for SPCA2C and suggests it may be involved in affecting CCDC47, STIM1 and Orai1 function.

## Introduction

Calcium (Ca^2+^) is an essential second messenger in multicompartment cells, aiding in the communication between organelles or with the extracellular environment (1). Signaling proteins and transcription factors are often regulated by Ca^2+^ and, therefore, changes in cytosolic Ca^2+^ influence a multitude of cellular processes including survival, proliferation, and gene expression (2–4). In acinar cells of the exocrine pancreas, control of spatial and temporal cytosolic Ca^2+^ levels is essential for regulated exocytosis of digestive enzymes (5, 6). Since aberrant Ca^2+^ signaling is associated with increased susceptibility and severity of pancreatic disease, understanding the interconnected pathways involved in intracellular Ca^2+^ regulation is critical (7).

One of the main pathways for Ca^2+^ influx in non-excitable cells, including pancreatic acinar cells, is store-operated Ca^2+^-entry (SOCE) (5). SOCE functions to replenish endoplasmic reticulum (ER) Ca^2+^ concentrations following inositol-trisphosphate receptor (IP_3_R)-mediated Ca^2+^ release from the ER and sustains high cytosolic Ca^2+^ levels to promote intracellular signaling (8, 9). While interactions between stromal interaction molecule 1 (STIM1) and the Ca^2+^ channel subunit Orai1 at ER-plasma membrane (PM) junctions result in an influx of Ca^2+^ into the cell following SOCE activation, many other proteins modulate this pathway including store-operated Ca^2+^ entry associated regulatory factor and transient receptor potential cation channel 1 (10–14). Orai1 can also promote changes in Ca^2+^ levels independent of ER Ca^2+^ concentrations through store-independent Ca^2+^ entry (SICE) (15). Small conductance Ca^2+^-activated K^+^ channel, K_V_10.1 and the secretory pathway Ca^2+^ ATPase 2 (SPCA2) interact with Orai1 to allow constitutive Ca^2+^ influx (15–17). However, only a C-terminal isoform for SPCA2 (SPCA2C) containing the last four exons of full-length SPCA2 is expressed to detectable levels in the pancreas (18, 19).

Like SPCA2, SPCA2C interacts with Orai1 to affect SICE (15, 20). Conversely, while SPCA2 is classified as a Golgi-resident Ca^2+^ ATPase, SPCA2C predominantly localizes to the ER in acinar cells (20, 21). Overexpression of SPCA2C in HEK293 cells increases SOCE, resting cytosolic and ER Ca^2+^ (20). Since SPCA2C is the only SPCA2 isoform expressed in pancreatic acinar cells and does not have a Ca^2+^ ATPase function to directly modulate cytosolic Ca^2+^ levels, we suggest SPCA2C regulates several pathways involved in Ca^2+^ homeostasis (19). The goal of this study was to define additional SPCA2C binding partners to understand its mechanistic role in Ca^2+^ regulation and acinar physiology.

To determine the SPCA2C interactome, we performed an unbiased screen in HEK-Orai1^YFP^ cells using a proximity-dependent biotin identification (BioID) system. Using a fusion protein with the BirA* enzyme attached to the C-terminus of SPCA2C, 123 candidate SPCA2C interactors were identified, with an additional 27 proteins revealed by Significance Analysis of INTeractome (SAINT) analysis. Gene Ontology (GO) and KEGG pathway analyses of these proteins supported SPCA2C’s localization to the ER and a role in Ca^2+^ regulation, but also uncovered potentially novel functions in vesicular transport. We identified CCDC47 as a high confidence SPCA2C-interacting protein involved in Ca^2+^ regulation. Interactions and co-localization between SPCA2C and CCDC47 were confirmed in HEK293 cells and included interactions with STIM1 and Orai1, supporting roles for both proteins in SOCE regulation. Overall, this study revealed a network of Ca^2+^ regulating proteins interacting with SPCA2C and provided insight into the molecular determinants of SPCA2C/CCDC47 interactions with STIM1 and Orai1. We propose SPCA2C and CCDC47 form a complex with STIM1 and Orai1 to modulate SOCE.

## Results

### In-silico modeling indicates the C-terminus of SPCA2C is unstable and orientated in the cytosol

No three-dimensional structure for SPCA2C has been experimentally elucidated to date. Thus, to gain insight into the topology of SPCA2C and guide the design of our fusion construct with BirA*, we generated an in-silico model of SPCA2C. We used MODELLER and the human SERCA2b pump cryo-electron microscopy (cryo-EM) structure (6LLE) as a template to generate a human SPCA2C structural model. Human SPCA2C was modeled since it showed the highest sequence similarity (i.e. 22 % identity and 62 % similarity over a 109 amino acid overlap) to any experimentally determined P-type ATPase structure (i.e. human SERCA2b). The resultant SPCA2C model revealed four transmembrane (TM) helices oriented similar to TM 7-10 of SERCA2b. A short non-TM helix was also predicted preceding the final TM (Figure 1A). We denote the five helices of our SPCA2C model as α1-α5, where α1, α2, α3 and α5 are equivalent to TM 7-10 of SERCA2b. We also generated an AlphaFold2 model of human SPCA2C, finding a highly similar five-helix predicted structure as generated with MODELLER [i.e. backbone root mean square deviation (RMSD) excluding the loops = 1.4 Å].

**Figure 1.**
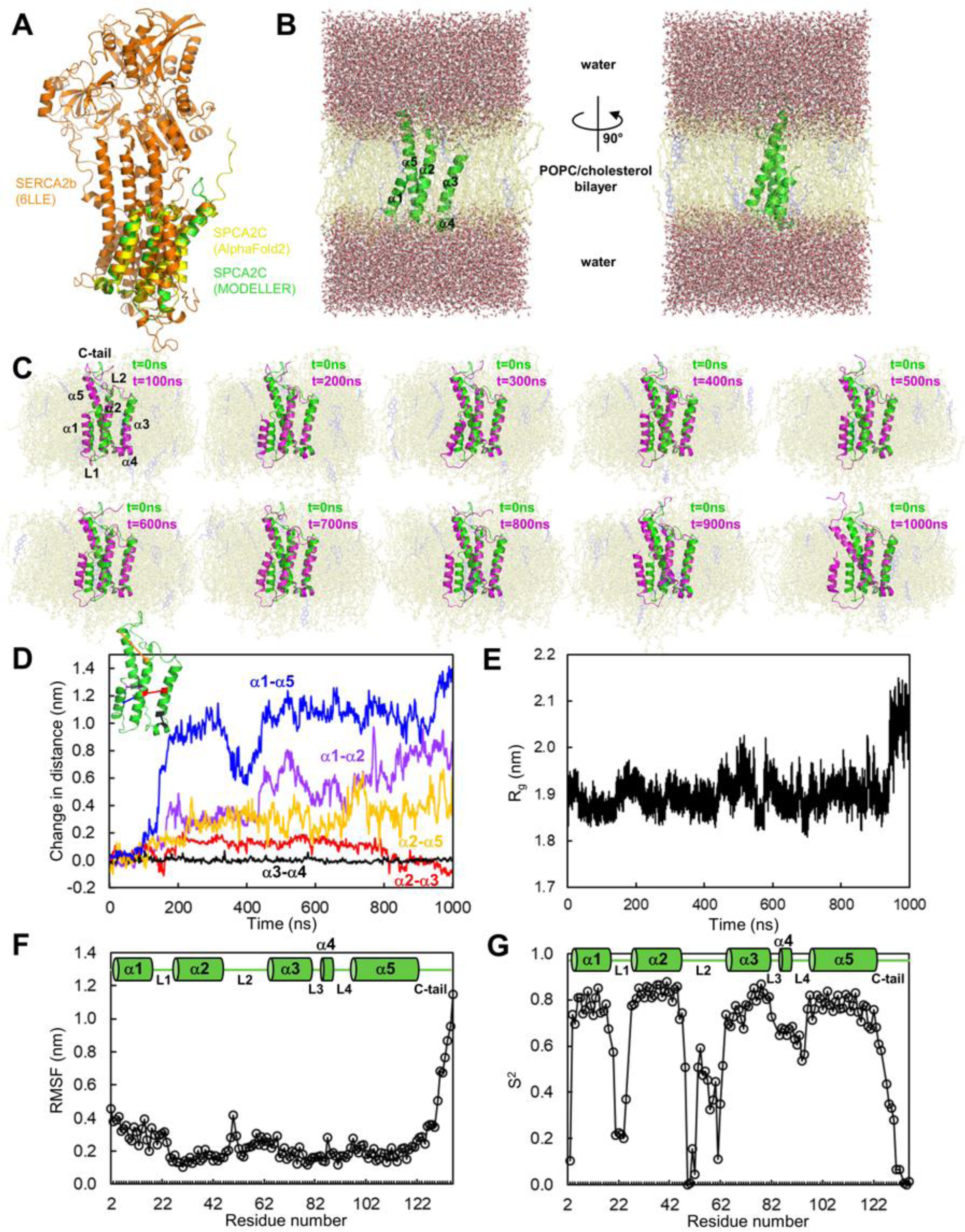
In silico modeling indicates SPCA2C is stable in a phospholipid bilayer with the C terminal extended into the cytoplasm. (**A**) Structural superposition of SERCA2b (6LLE), human SPCA2C modeled using AlphaFold2 and human SPCA2C generated using MODELLER (orange, yellow and green backbone cartoon representation, respectively). (**B**) Complete system generated using the CHARMM-GUI for MD simulations. Human SPCA2C (green backbone) is centered in a POPC (pale yellow sticks)/cholesterol (light blue sticks) bilayer. The bilayer is sandwiched between 30 Å thick water layers (red and white ball and sticks). (**C**) Conformational changes observed for human SPCA2C over the 1000 ns simulation time. In each still image, the SPCA2C conformation is shown at the specified time point (magenta backbone) relative to the starting conformation (green backbone). Only the bilayer (transparent pale yellow/light blue sticks) and protein are shown for clarity. (**D**) Change in helix-helix distances over the 1000 ns simulation time. The α1-α5 distances (blue) were measured from F12 (α1) to F101 (α5); the α1-α2 distances (purple) were measured from I5 (α1) to F39 (α2); the α2-α5 distances (orange) were measured using C47 (α2) to C123 (α5); the α2-α3 distances (red) were measured using V37 (α2) to S72 (α3); the α3-α4 distances (black) were measured using V79 (α3) to L85 (α4). All distances were measured between Cα atoms, and changes were determined relative to the t=0 ns conformation. The inset shows the arbitrarily chosen positions on the SPCA2C structure (green backbone) for the measurements, colour-coded to match the plot. (**E**) Radius of gyration (Rg) of SPCA2C over the 1000 ns simulation time. (**F**) Root mean square fluctuation (RMSF) of each residue during the 1000 ns. RMSF was calculated relative to the t=0 ns conformation. (G) Order parameters (S2) of each backbone amide during the 1000 ns simulation time. In F and G, the secondary structure components (green) relative to the residue number are shown above the plots, with α-helices represented by cylinders and loops by straight lines.

Since SPCA2C is transcribed only from the last four exons of *ATP2C2* and has fewer domains than P-type ATPases, we next used GROMACS molecular dynamics (MD) simulations to assess the structural and topological stability of our SPCA2C model in a 1-palmytoyl-2-oleoyl-sn-glycero-3-phosphocholine (POPC)/cholesterol bilayer. The bilayer was surrounded by a 30 Å water shell (Figure 1B). Over a 1000 ns simulation time, structural changes were predominantly localized to two motifs: the amino (N)-terminal α1 and the carboxyl (C)-terminal α5. In contrast, the position and conformation of α2, α3 and α4 remained relatively stable (Figure 1C; Movie S1). After the first ∼100 ns, α1 showed progressively increased distance from the stable α2/α3/α4 cluster over the 1000 ns. Similarly, the C-terminal portion of α5, exhibited increased distance from the central α2. Quantitatively, the α1-α2 distance increased by ∼8 Å and the α5-α2 distance increased by ∼5 Å, over the simulation time. These helical movements resulted in an increased α1-α5 distance of ∼14 Å over the 1000 ns. In contrast, the α3-α2 and α3-α4 change in distances rarely exceeded ∼1.5 Å over the simulation time (Figure 1D).

Despite the notable repositioning of α1 and α5 over the 1000 ns, the radius of gyration (R_g_) of the protein did not appreciably change, arguing against global unfolding on the ns-µs timescale (Figure 1E). We calculated the root mean square fluctuation (RMSF) on a per residue basis, finding RMSF was strikingly higher for residues of the C-terminus compared to all other residues (Figure 1F). This RMSF calculation provides information on the distance fluctuation of each residue relative to the average position over the simulation time. The order parameters (S2) for each backbone amide approached 0 in loop1, loop2 and the C-terminal region (Figure 1G). The S2 calculation measures the flexibility of each backbone amide (N-H) bond vector with values of 0 and 1 indicating complete flexibility and rigidity, respectively, on a ps timescale. Thus, the C-terminal region of SPCA2C shows the most flexible residues on the ns timescale, while loop1, loop2 and the C-terminal region showed the highest and similarly dynamic backbone amides on a ps timescale. The separability of the timescale of these dynamics is evident in the N-terminal end of α3, which showed high mobility in the ps but not ns timescales (Figure 1F and 1G).

Collectively, our modeling and MD simulations suggest SPCA2C adopts a conformation similar to the TM 7-10 region of SERCA2b, with the C-terminal tail localized to the cytosol. While the overall fold remained relatively intact over the ns-µs timescale, the C-terminal tail was the only region that was highly dynamic in both ps and ns timescales. Given that flexible regions of protein are often the sites of protein interaction and the C-terminal region of SPCA2C was the most dynamic on the ps and ns timescales, we fused BirA* to the SPCA2C C-terminal tail to minimize distances between the biotinylating enzyme and proteins contacting the C-terminal region, avoid destabilizing the helices, and avert influencing the mobility of the SPCA2C helices.

### BioID assay identifies 150 putative SPCA2C- interacting proteins with ER and exocrine-related function

To determine the SPCA2C interactome, HEK-Orai1^YFP^ cells transiently expressing SPCA2C-BirA^*HA^ were incubated with biotin for 24 hours, allowing labeling of any proteins within approximately 10 nm of SPCA2C (22). Fusion of the 35 kDa BirA* enzyme did not alter localization of SPCA2C within the cell as SPCA2C-BirA^*HA^ overlapped with SPCA2C^mRFP^ (Figure S1A) and showed areas of overlap with Orai1^YFP^ in HEK293 cells, as observed previously (Figure S1B) (20). Western blot analysis of biotinylated proteins detected several distinct bands in membrane cell fractions where SPCA2C-BirA^*HA^ expression was also observed (∼35 kDa; Figure 2A) while only two bands were evident in control samples with BirA^*HA^ expression. SPCA2C-BirA^*HA^ co-immunoprecipitated with Orai1, further confirming fusion of the BirA* enzyme to the C-terminus of SPCA2C does not affect localization or disrupt interactions with other proteins. Visualization of biotinylated proteins with Streptavidin-Alexa Flour 568 showed overlap with SPCA2C-BirA^*HA^ (Figure S1C). This localization was distinct from biotinylated proteins in BirA*-expressing cells, indicating the biotinylation of proteins was specific to SPCA2C localization.

**Figure 2.**
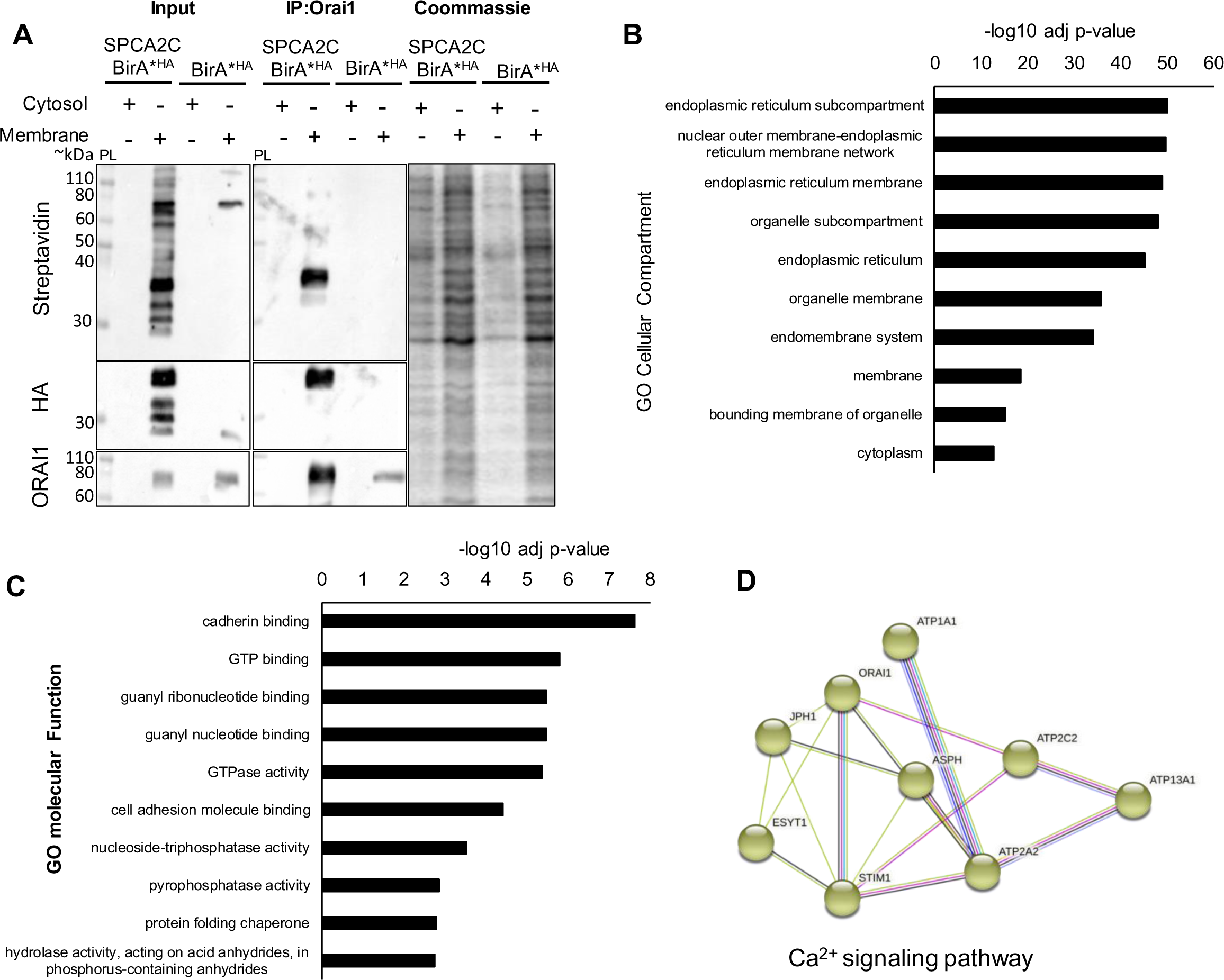
Bioinformatic analysis of proteins identified from BIOID screen. (**A**) Co-IP analysis examining the expression of SPCA2C-BirA^*HA^, Orai1^YFP^ and biotin-labeled proteins detected with Streptavidin-HRP, in Input and following IP of Orai1^YFP^. Cytosol and membrane fractions of protein were collected from HEK-Orai1^YFP^ cells transiently expressing SPCA2C-BirA^*HA^ or BirA^*HA^ (negative control) after incubation with biotin for 24 hours. Input shows multiple bands of biotinylated proteins in membrane fraction of SPCA2C-BirA^*HA^ expressing cells and only a few bands in membrane fraction from BirA^*HA^ expressing cells. As a positive control, co-IP of SPCA2C-BirA^*HA^ with Orai1^YFP^ was observed. Gel stained with Coomassie blue shows relative protein loading. PL= protein ladder. The top ten GO terms of high-confidence interactors were found by BIOID assay for (**B**) cellular compartments and (**C**) molecular function. (**D**) A protein-protein interaction network of high-confidence interactors was established by STRING and groups of proteins were clustered by the ClusterOne plug-in in Cytoscape. Groups were classified based on GO Biological Process, Molecular Function, or KEGG pathway analysis of collective proteins in each group.

Proteins biotinylated by SPCA2C-BirA^*HA^ were identified via streptavidin affinity (SA) capture followed by mass spectrometry (MS) analysis. Whole protein lysates were collected rather than membrane-enriched fractions to ensure all biotinylated proteins were represented in the SA-purified MS analysis regardless of where they localized. To develop a list of proteins most likely to interact with SPCA2C, proteins detected by MS following SA-precipitation from SPCA2C-BirA^*HA^-expressing cells were referenced against proteins detected in the empty control BirA^*HA^ SA-precipitation (Supplemental File 1). Proteins listed in both SPCA2C-BirA*^HA^ and BirA^*HA^ MS lists or not in all SPCA2C-BirA^*HA^ replicates were omitted from further analysis unless they received a SAINT score greater than or equal to 0.95 (27 proteins), suggesting a true interactor (Figure S2). In total, 150 proteins were identified as potential members of the SPCA2C interactome based on the unique identification in all SPCA2C-BirA^*HA^ replicates and SAINT score. This list included Orai1, unique to SPCA2C-BirA^*HA^ replicates, confirming the screen identified known interacting partners for SPCA2C.

Gene Ontology (GO) analysis using the 150 putative interacting proteins identified multiple terms linked to the ER membrane and ER-related molecular processes, including “protein folding chaperone”, supporting previous IF analysis of SPCA2C localization (Figure 2B, C) (20). To gain a better understanding of the functional complexes represented in the SPCA2C interactome, the interactive network of proteins was visualized using STRING analysis and Cytoscape. Using ClusterOne in Cytoscape, the network was arranged by distributing highly interconnected proteins into groups and a total of 23 groups were identified. These groups are likely to be multi-protein complexes, of two or more proteins and were further labeled with functional terms from GO or KEGG pathway analysis. Several groups contained clusters suggesting function and localization to the ER, while additional pathways include those involved in vesicle-mediated trafficking, protein processing and protein export, suggesting novel roles for SPCA2C associated with acinar cell physiology (Figure S3). A network of proteins involved in Ca^2+^ signaling was also identified, supporting previous findings SPCA2C aids in maintaining cytosolic and ER Ca^2+^ homeostasis (Figure 2D).

### Confirmed interactions between SPCA2C and Ca^2+^ signaling proteins from BioID screen

Despite SPCA2C lacking an ATPase domain required for active Ca^2+^ transport, our previous studies showed SPCA2C influenced cellular Ca^2+^ homeostasis through alternate pathways, including SICE and SOCE (19, 20). Therefore, we verified proteins within the SPCA2C interactome clustered into the Ca^2+^ signaling pathway. Biotinylated forms of ESYT1, STIM1, and Orai1 were all detected by SA pull-down in whole cell lysates from HEK-Orai1^YFP^ SPCA2C-BirA^*HA^-expressing cells incubated 24 hours with biotin (Figure 3A). Interestingly, analysis of ASPH pull-down with Streptavidin resulted in detection of an ASPH isoform consistent with the molecular weight of junctate (∼50 kDa; Figure 3A) (23). We observed co-IP of ESYT1 with SPCA2C-BirA^*HA^ (Figure 3B), although this interaction was verified ∼50% of the time, suggesting a weak or transient interaction between the two proteins. Reverse co-IP for ASPH confirmed interactions with SPCA2C-BirA^*HA^ (Figure 3C). We also observed interaction between STIM1 and SPCA2C^FLAG^, through co-IP for FLAG or reverse co-IP with an antibody for STIM1 (Figure 3D). Overall, these findings supported our BioID screen for the SPCA2C interactome and identified previously unknown interactions between SPCA2C and ASPH, ESYT1, and STIM1.

**Figure 3.**
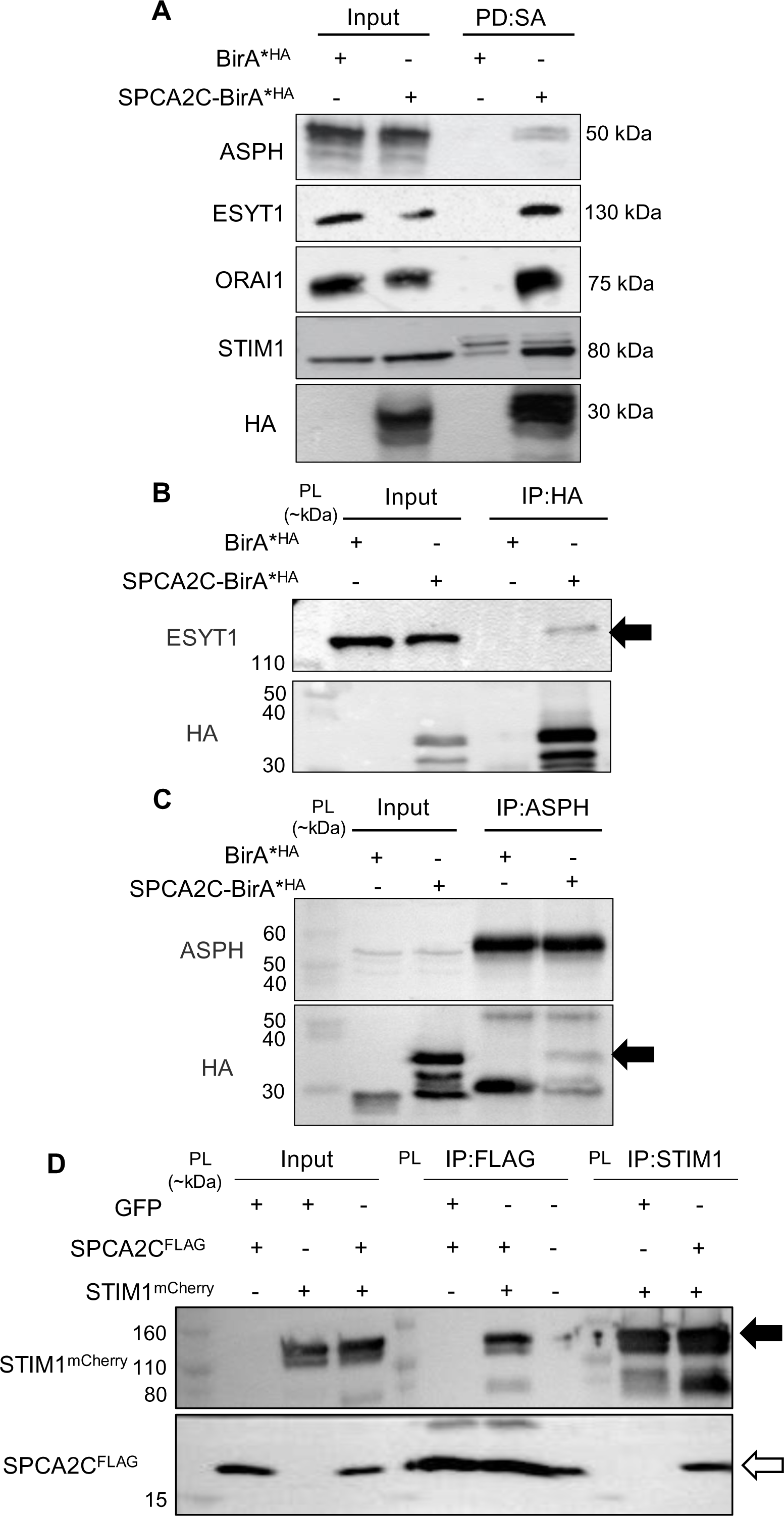
Confirmation of SPCA2C interactions with proteins identified from BIOID screen involved in Ca^2+^ regulation. (**A**) Streptavidin (SA) pull-down from whole cell lysates of biotin-treated HEK-Orai1^YFP^ cells with exogenous SPCA2C-BirA^*HA^ or BirA^*HA^ expression. In samples with SPCA2C-BirA^*HA^ expression, western blot detected pull-down of proteins identified from the screen that clustered in the Ca^2+^ signaling pathway (ASPH, ESYT1, Orai1, and STIM1). (**B-D**) Co-IP analyses confirming interactions between SPCA2C and the identified subset of proteins. (**B**) Co-IP of endogenous ESYT1 (black arrow) with SPCA2C-BirA^*HA^, and (**C**) SPCA2C-BirA^*HA^ (black arrow) with endogenous ASPH. (**D**) Co-IP of STIM1^mCherry^ (black arrow) with SPCA2C^FLAG^ (white arrow) and reverse co-IP of STIM1^mCherry^ with SPCA2C^FLAG^. All co-IPs performed with whole cell lysates from HEK293 cells. PL= protein ladder.

### SPCA2C co-localizes and interacts with CCDC47

To provide more detail on a putative interacting partner for SPCA2C, we focused on coiled-coil domain-containing protein 47 (CCDC47; also referred to as calumin). CCDC47 was chosen since it was ranked as the highest confidence interacting protein with a known function in Ca^2+^ homeostasis (FC score= 25.62, SAINT score=1; Table 1) (24). While STRING and ClusterOne analyses did not group CCDC47 with Ca^2+^ signaling proteins (Figure S3), previous studies showed CCDC47 interactions with Orai1 and STIM1 and overexpression of CCDC47 increased both ER Ca^2+^ levels and SOCE responses (25, 26). Interaction between SPCA2C and CCDC47 was confirmed through co-IP of SPCA2C-BirA^*HA^ with endogenous CCDC47 in HEK293 cells and reverse co-IP using a CCDC47-specific antibody (Figure 4A, B). Interaction between the two proteins was observed in the absence of the BirA* enzyme as transiently transfected SPCA2C^FLAG^ co-immunoprecipitated with CCDC47 (Figure 4C). Spinning disk confocal microscopy and line scan analysis revealed co-localization between SPCA2C^mRFP^ and IF-stained CCDC47^FLAG^ in HEK293 cells (Figure 4D, E) confirming interaction and co-localization between SPCA2C and CCDC47.

**Table 1:** Proteins within the SPCA2C interactome based on SAINT analysis and related to Ca^2+^.

| Protein ID | Gene | Max FCA | Function | Reference |
| --- | --- | --- | --- | --- |
| CCD47 | *CCDC47* | 25.62 | Regulates Ca<sup>2+</sup> homeostasis in the endoplasmic reticulum | Zhang et al (2007) *Cell Calcium* |
| VAPB | *VAPB* | 19.78 | Recruits TRPC3 to ER/PM junction and cellular Ca<sup>2+</sup> homeostasis | Liu et al (2022) *JCB*<br>Ma et al (2023) *J Mol Cell Biol* |
| ITB1 | *ITGB1* | 15.84 | Expression increased by Ca<sup>2+</sup>, related to calpain 2 | Li et al (2019) *Cell Signal* |
| CALX | *CANX* | 10.92 | Protein chaperone similar to Calreticulin | Paskevicius et al (2023) *Cells* |
| STIM1 | *STIM1* | 9.63 | Involved in Store-operated Ca<sup>2+</sup> entry |  |
| NB5R1 | *CYB5R1* | 8.43 | Associated with Ca<sup>2+</sup> pumps in lipid rafts-associated nanodomains | Marques-da-Silva and Gutierrez-Merino (2014) *Cell Calcium* |
| ASPH | *ASPH* | 8.19 | Structural component where ORAI1 and STIM1 localize | Srikanth et al (2012) *PNAS* |
| ESYT1 | *ESYT1* | 6.15 | Binds Ca<sup>2+</sup> and translocates to ER-PM sites in response to increased cytosolic Ca<sup>2+</sup> levels |  |
| CLGN | *CLGN* | 5.51 | Protein chaperone similar to Calreticulin |  |
| PROF1 | *PFN1* | 3.28 | Related to Ca<sup>2+</sup> signaling in bladder cancer | Frantzi et al (2016) *Oncotarget* |
| AT2A2 | *ATP2A2* | 3.78 | Ca<sup>2+</sup> ATPase related to Darier disease | Savignac et al (2011) *BBA* |
| RCN1 | *RCN1* | 3.76 | Interacts with IP3 receptors and inhibits Ca<sup>2+</sup> release | Xu et al (2017) *Oncogenesis* |
| TAGL2 | *TAGLN2* | 2.7 | Belongs to the family of calponin proteins involved in Ca<sup>2+</sup> mobilization | Yim et al (2020) *Front Cell Dev Biol* |

<sup>1</sup>Ranks given from 108 proteins based on FC score; <sup>2</sup>Fold change (FC): Based on the ratio of average normalized spectral counts in bait purifications to negative controls. Higher the number indicates a more likely interaction; <sup>3</sup>Significance Analysis of Interactome (SAINT): Provides a statistical conversion of the FC score onto the probability scale using the mixed-model analysis of underlying spectral count distribution; <sup>4</sup>Function based on literature review. Gray rows indicate those proteins identified through SAINT but not in all three BioID screens.

**Figure 4.**
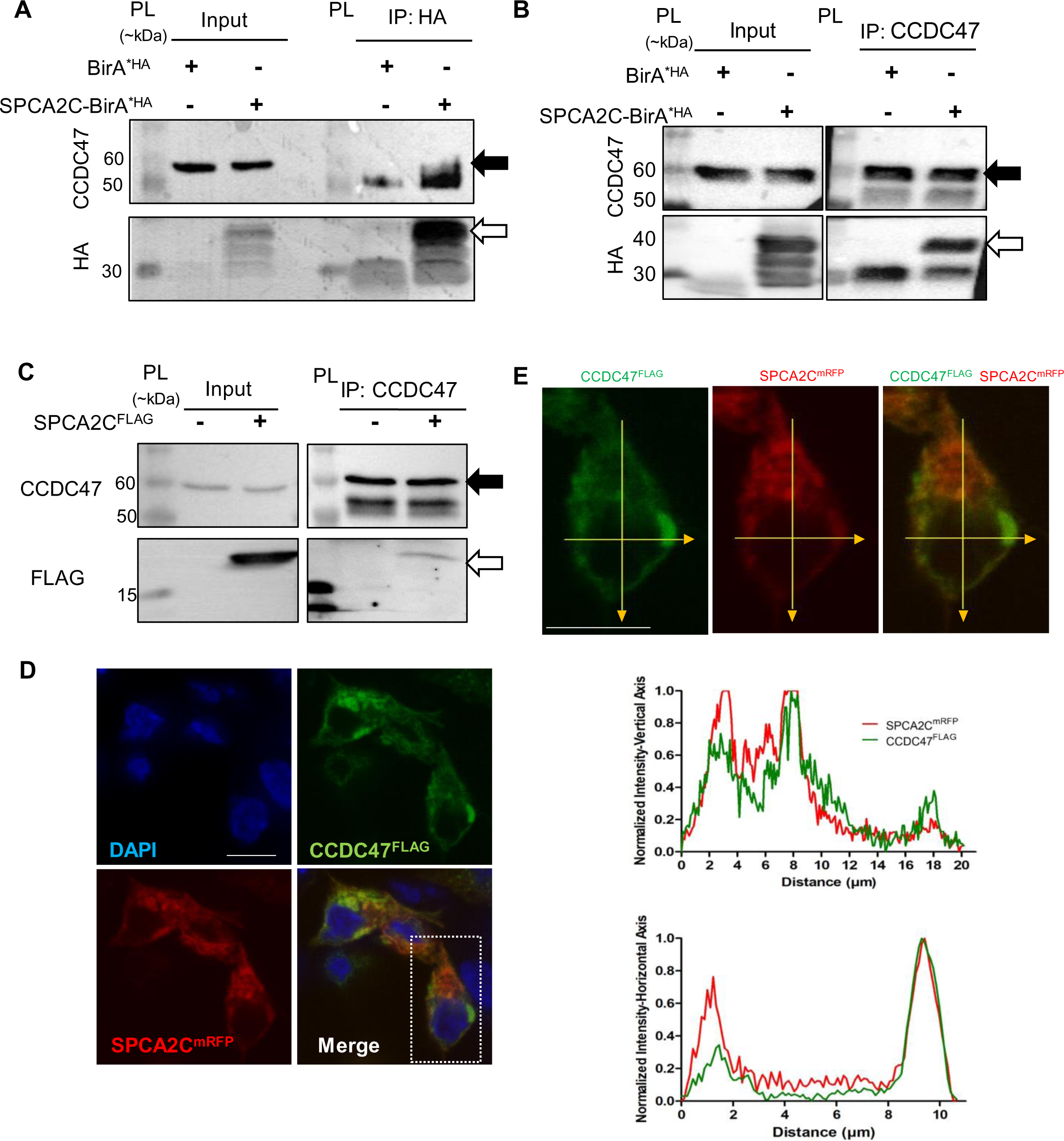
Interactions and co-localization between SPCA2C and CCDC47. (**A-C**) Co-IP analyses of SPCA2C-CCDC47 interactions in HEK293 cells. (**A**) Co-IP of endogenous CCDC47 with SPCA2C-BirA^*HA^ and (**B**) reverse co-IP of SPCA2C-BirA^*HA^ with CCDC47. IPs performed with BirA^*HA^ only are negative controls. (**C**) Co-IP of SPCA2C^FLAG^ with CCDC47. Black arrows indicate CCDC47 expression. White arrows indicate SPCA2C-BirA^*HA^ or SPCA2C^FLAG^ expression. PL=protein ladder. (**D**) Single plane confocal image (0.3 µm depth) of SPCA2C^mRFP^ and IF-stained CCDC47^FLAG^. White arrows indicate areas of colocalization. Nuclei stained with DAPI. Boxed area indicates cells used for line scan analysis in (**E**). Normalized fluorescent intensities of pixels along horizontal and vertical yellow lines were plotted for SPCA2C^mRFP^ and fluorescein-stained CCDC47^FLAG^ as a function of distance (µm). Scale bars= 10 µm.

To identify the regions within CCDC47 required for SPCA2C interaction and determine whether these regions were involved also in interactions with STIM1 and Orai1, we performed deletion analysis for CCDC47. Since CCDC47 is an ER-resident protein consisting of a luminal domain, single-pass transmembrane (TM) domain, and a cytosolic domain containing α-helical globular and coiled-coil regions (24), we decided to delete these regions. Full-length CCDC47 (CCDC47_full_) or CCDC47 truncations were subcloned into a *pDsRed2-ER* mammalian expression vector. Truncations targeted the cytosolic coiled-coil domain (CCDC47_ΔCC_; deletion of residues 400-483), the cytosolic globular and coiled-coil domains (CCDC47_ΔG+ΔCC_; deletion of residues 161-483), or the luminal and cytosolic domains (CCDC47_TM_; deletion of residues 1-130 and 161-483, respectively, leaving only the TM region). Full-length and CCDC47 truncations lacked residues 1-20 at the N-terminus, which correspond to the ER signal sequence since *pDsRed2-ER* encodes an ER localization sequence, thereby maintaining ER targeting for these constructs. Following co-expression of the CCDC47 truncations with SPCA2C^FLAG^ in HEK293 cells, SPCA2C^FLAG^ was pulled down with all CCDC47 proteins, suggesting interactions with SPCA2C occur via the transmembrane domain of CCDC47 (Figure 5B). Conversely, co-IP of STIM1^mCherry^ with the CCDC47 truncations identified regions outside the TM of CCDC47 that were important for interaction (Figure 5C). IP for the MYC-tagged CCDC47 constructs showed reduced interaction between CCDC47 and STIM1 only when the coiled-coil domain of CCDC47 was deleted (CCDC47_ΔCC_) while deletion of the entire cytosolic domain, including both globular and coiled-coil domains (CCDC47_ΔG+ΔCC_), restored interaction. These results suggest the coiled-coiled domain stabilizes the CCDC47-STIM1 interaction and the presence of the globular domain in the absence of the coiled-coiled domain promotes de-stabilization of that interaction. Similarly, Orai1 interacted with full-length CCDC47_full_, while CCDC47_ΔG+ΔCC,_ and CCDC47_TM_ showed reduced interaction, and CCDC47_ΔCC_ showed no interaction (Figure 5D). These results collectively suggest the transmembrane domain of CCDC47 is sufficient for interaction with SPCA2C, STIM1, and Orai1, and deletion of the coiled-coil domain in the presence of the globular domain results in weakened interactions with STIM1 and Orai1.

**Figure 5.**
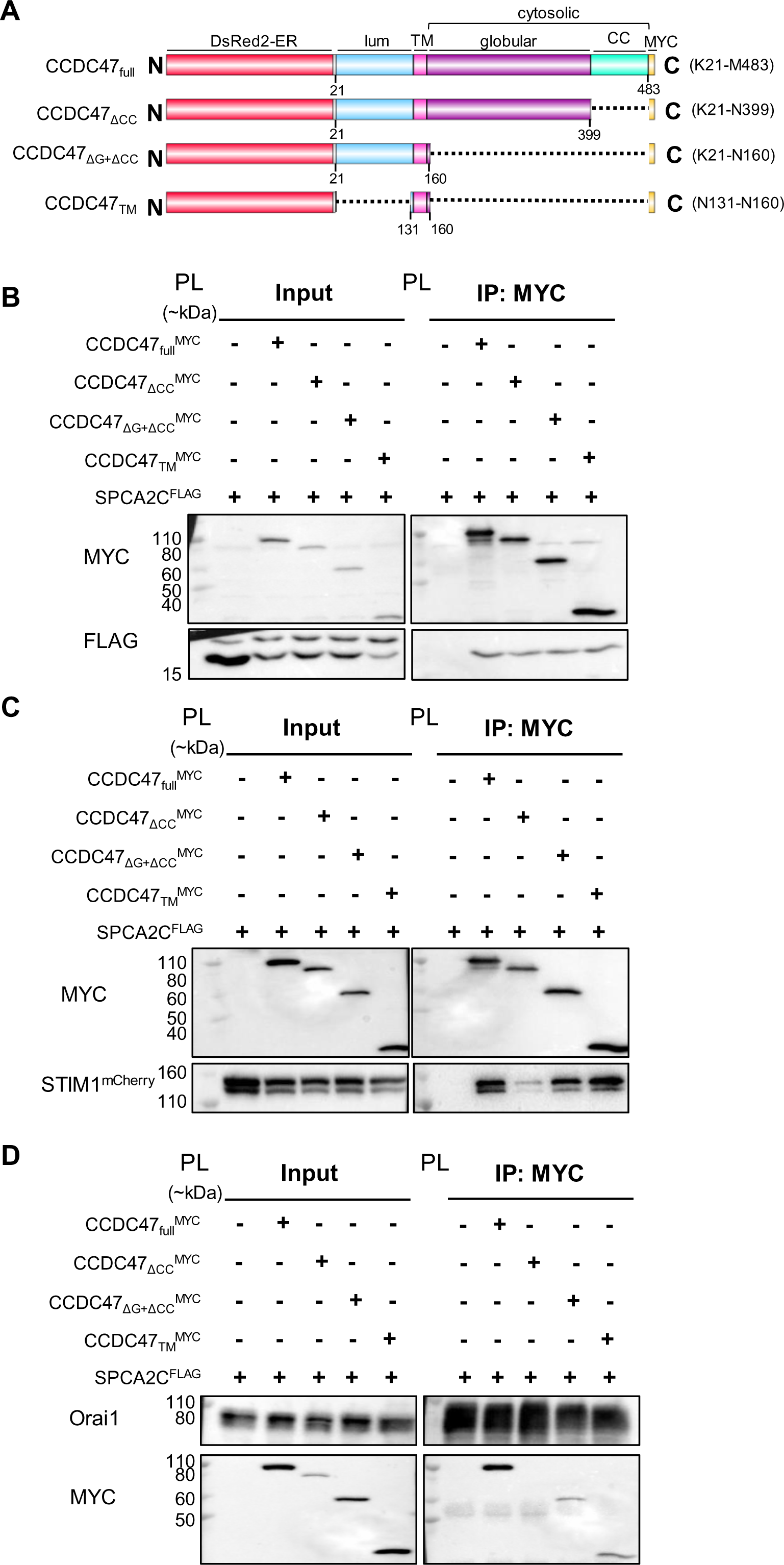
Interactions between CCDC47 truncations and SPCA2C, Orai1, and STIM1. (**A**) Domain architecture diagram of DsRed2 fused to MYC-tagged CCDC47 truncations. Full-length CCDC47 contains an ER luminal domain (lum), single-pass transmembrane domain (TM), and a cytosolic domain with an alpha-helical globular domain (G) and coiled-coil (CC) domain. Full-length CCDC47 (CCDC47_full_), CCDC47 with deletion of coiled-coil domain (CCDC47_ΔCC_), deletion of both globular and coiled-coil domains (CCDC47_ΔG+ΔCC_), and deletion of both luminal and cytosolic domains (CCDC47_TM_) were examined. Dotted line represents deleted domains and the specific residues spanning each recombinant CCDC47 protein are indicated in brackets. (**B**) Expression of SPCA2C^FLAG^ detected following immunoprecipitation of MYC tagged DsRed2-CCDC47_full_, -CCDC47_ΔCC_, – CCDC47_ΔG+ΔCC_, and – CCDC47_TM_. (**C**) Co-IP of STIM1^mCherry^ with DsRed2-CCDC47 ^MYC^, -CCDC47 ^MYC^, – CCDC47 ^MYC^ and – CCDC47 ^MYC^. (**D**) Co-IP of DsRed2-CCDC47 ^MYC^, -CCDC47 ^MYC^, – CCDC47 ^MYC^ and – CCDC47 ^MYC^ with Orai1^YFP^. Co-IPs were performed using cell lysates from transiently transfected HEK293 or HEK-Orai1^YFP^ cells. PL=protein ladder. Schematic created using DOG (Domain Graph, version 2.0) (47).

We next visualized co-localization of DsRed2-fused CCDC47 truncations and SPCA2C, STIM1 or Orai1 to determine whether specific domains of CCDC47 influenced co-localization with SOCE-regulating proteins. Fluorescent analysis for DsRed2 localization indicated deletion of the coiled-coil domain of CCDC47 (CCDC47_ΔCC_) resulted in increased plaque formation compared to the CCDC47_full_, CCDC47_ΔG+ΔCC,_ and CCDC47_TM_ constructs. Colocalization with SPCA2C was maintained for each CCDC47 truncation (Figure 6A) and the increased plaque formation for CCDC47_ΔCC_ was observed regardless of whether SPCA2C, STIM1 or Orai1 were co-expressed (Figure S4). Interestingly, SPCA2C localized to these plaques more consistently than STIM1 and Orai1 consistent with co-IP data suggesting that reduced CCDC47-STIM1 and -Orai1 interactions in the absence of the coiled-coiled domain may be due to altered localization. Colocalization of all CCDC47 truncations with STIM1 and Orai1 was observed (Figure 6B,C), but there was diminished localization between CCDC47_TM_ and STIM1 (Figure 6B). Overall, the changes in localization of CCDC47 truncations resulted in more consistent colocalization with SPCA2C compared to STIM1 and Orai1.

**Figure 6.**
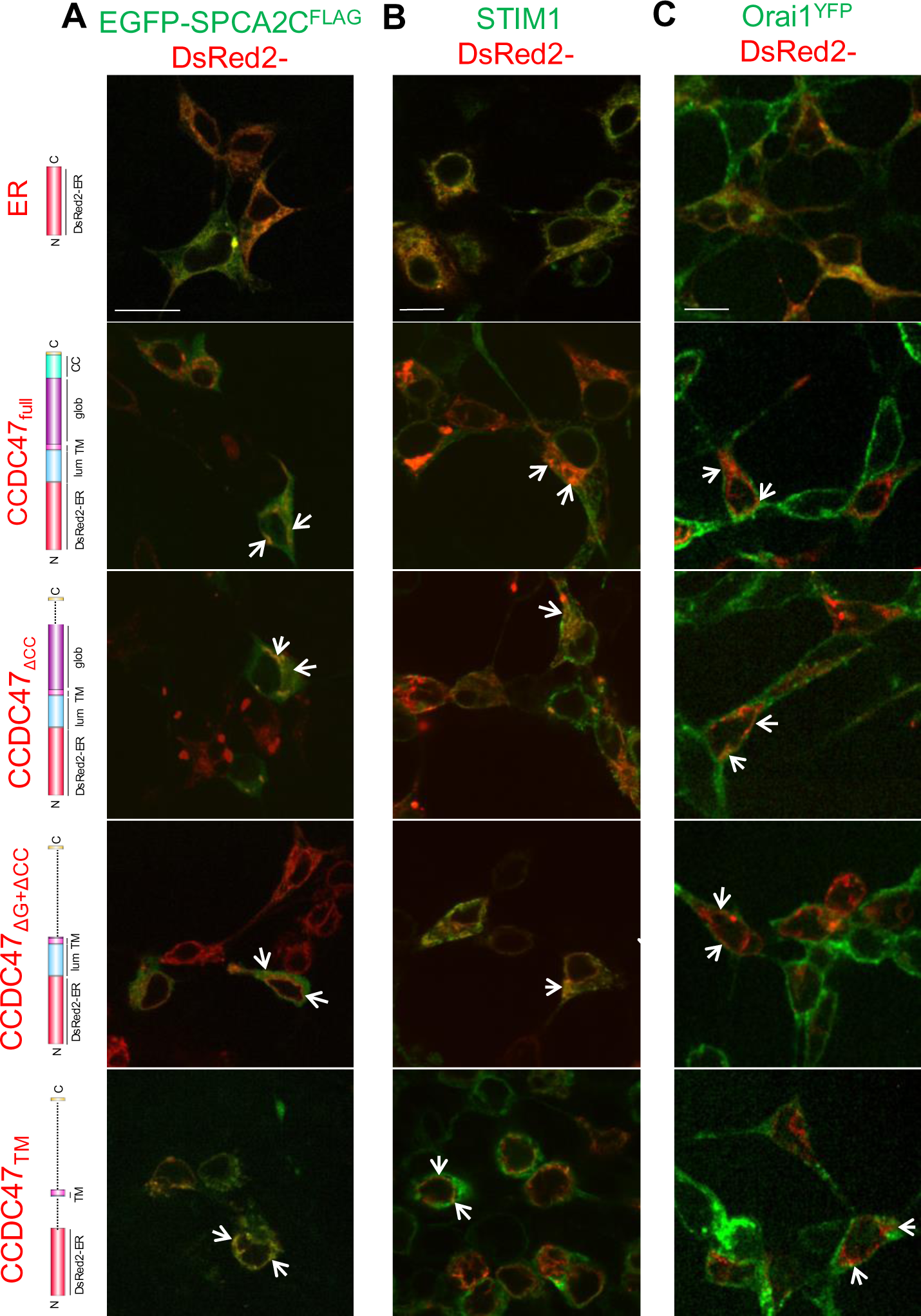
Colocalization of CCDC47 truncations with SPCA2C, STIM1 or Orai1. Merged spinning disk confocal microscopy images of DsRed2 fused to CCDC47 truncations (red) with (**A**) EGFP-SPCA2C^FLAG^, (**B**) immunofluorescent stained STIM1, or (**C**) Orai1^YFP^ (green). Truncations of CCDC47 are delineated in domain architecture diagrams on the left and include CCDC47_full_, CCDC47_ΔCC_, CCDC47_ΔG+ΔC_, and CCDC47_TM_. Visualization of the DsRed2-ER protein without fusion to CCDC47 was used as a control to ensure CCDC47-specific localization. Areas of colocalization of CCDC47 constructs with SPCA2C, STIM1, or Orai1 are indicated with white arrows. Images show single-plane images at 0.3 µm thickness. Each scale bar represents 10 µm and applies to images in the corresponding column. Imaging performed in HEK293 or HEK-Orai1^YFP^ cells.

## Discussion

The regulation of Ca^2+^ is critical to the function and survival of eukaryotic cells and altered Ca^2+^ signaling underlies numerous disease processes (27, 28). Despite the absence of domains required for binding or transporting Ca^2+^ as a P-type ATPase, SPCA2C has a unique function in Ca^2+^ homeostasis, affecting multiple cellular pathways mediating or regulating Ca^2+^ movement (20). In this study, we identified SPCA2C interactions with several proteins involved in SOCE. While interactions with Orai1 have been documented, we identified previously unknown interactions between SPCA2C and STIM1 and CCDC47, suggesting a regulatory role for SPCA2C in SOCE. We also identified additional protein interactors that suggests SPCA2C has functions outside of Ca^2+^ regulation and reflects a more central role in acinar cell biology.

### The SPCA2C interactome suggests additional non-Ca^2+^ signaling roles

Previous work suggested SPCA2C interacts with pathways involved in Ca^2+^ signaling but only interactions with Orai1 have been documented to date (20). Therefore, an unbiased screen was performed to identify other interacting proteins for SPCA2C. In total, 150 proteins were identified as part of the SPCA2C interactome and supported roles for SPCA2C in regulating cytosolic and ER Ca^2+^ homeostasis. This study is the first to identify interactions between SPCA2C and an extensive network of Ca^2+^ regulators, including CCDC47, STIM1, ESYT1, and ASPH. BioID was selected due to several advantages over other techniques such as yeast-two-hybrid (29). The BirA* enzyme quickly and irreversibly labels proteins, allowing the identification of transient or weak interactors that occur *in-cellulo*, which we suggest was instrumental in expanding the SPCA2C interactome. For example, while ESYT1 expression was detected following SA pull-down of biotinylated proteins, co-IP between SPCA2C and ESYT1 were inconsistent, suggesting weak or transient interactions.

This study also identified potentially unknown roles for SPCA2C in cell functions relevant to acinar cell biology. The most enriched pathway based on STRING analysis was vesicle-mediated trafficking, suggesting SPCA2C’s possible involvement in protein transport, an activity central to pancreatic acinar cells. Proenzyme storage and exocytosis require tightly regulated Ca^2+^ signaling, and previous analysis showed negligible SPCA2C expression in pancreatic acinar cells of mice lacking MIST1 (*Mist1^−/−^)* (18). Since *Mist1^−/−^* acinar cells show altered zymogen granule localization and decreased regulated exocytosis, this interactome data supports SPCA2C as an integral protein in the *Mist1^−/−^* phenotype, potentially coordinating both Ca^2+^ signaling and exocytosis (30). It is possible that the inclusion of vesicle-mediated trafficking proteins reflects the mechanism of SPCA2C’s translocation to its cellular localization as a limitation of the BioID assay is the identification of proteins in close proximity but not directly interacting. Future analysis of these putative interacting proteins are needed to define all of SPCA2C’s functions in pancreatic acinar cell physiology.

### SPCA2C interacts with multiple members of the SOCE pathway

Previous studies linked the full length SPCA2 to Store Independent Ca^2+^ entry (SICE) but suggested it does not regulate SOCE (15). Our SPCA2C-specific analysis indicates this is a distinguishing feature between the two isoforms (20). To date, we have only shown interaction with Orai1. However, STRING analysis of the SPCA2C interactome revealed several proteins with a function in Ca^2+^ regulation. We confirmed SPCA2C interactions with ESYT1 and ASPH, two proteins involved in mediating Ca^2+^ signals between the ER and outside the cell. ESYT1 translocates to ER-PM junctions in response to elevated cytosolic Ca^2+^ concentrations to increase ER-PM junctions and contributes to the enrichment of phosphatidylinositol 4,5-bisphosphate at the PM (31). ESYT1 is also involved in remodelling ER-PM contact sites into more stable ring-shaped structures, enhancing the replenishment of ER Ca^2+^ stores (32). ASPH hydroxylates asparagine and aspartic acid residues within epidermal growth factor-like protein domains and undergoes alternative splicing to produce multiple isoforms (33). We observed interaction of SPCA2C with an ASPH isoform consistent with the molecular weight of junctate, which interacts with the STIM1-Orai1 complex and aids in the recruitment of STIM1 to ER-PM junctions, increasing SOCE (23). This is also the first time interaction has been documented between SPCA2C and STIM1. While our previous work showed SPCA2C-Orai1 interactions were disrupted by overexpression of STIM1 (20), whether this was due to STIM1 displacing SPCA2C was unclear. In the current study, we show STIM1 interacts with SPCA2C in cells with little to no Orai1 expression, confirming SPCA2C interacts with several components of the SOCE complex. Since ESYT1 and junctate both have functions at ER-PM junctions, it is possible that SPCA2C may regulate STIM1 interactions with Orai1 at ER-PM contact sites (23, 31). Interaction with CCDC47 also support such a role.

We focused further characterization on CCDC47 since it was ranked as the highest confidence interacting protein related to Ca^2+^ signaling. Although bioinformatic analysis did not cluster CCDC47 with proteins involved in Ca^2+^ regulation, multiple studies show CCDC47 influences SOCE (24, 26, 34). *Ccdc47*^−/−^ MEFs and primary human dermal fibroblasts without CCDC47 expression demonstrate decreased ER Ca^2+^ levels and reduced SOCE compared to respective cells expressing CCDC47 (24, 34), while overexpression of CCDC47 in myocytes significantly increased ER Ca^2+^ stores and SOCE (26). Similar overexpression of SPCA2C increases ER Ca^2+^ and Ca^2+^ influx following store depletion (20). Since both SPCA2C and CCDC47 directly interact with Orai1 and STIM1, we suggest SPCA2C-CCDC47 interactions may regulate the STIM1-Orai1 complex and SOCE.

Using CCDC47 constructs with varying domain deletions, we showed the interaction between SPCA2C and CCDC47 occurs through the transmembrane domain of CCDC47. The α1 and α5 transmembrane domains of SPCA2C are most likely to interact with the CCDC47 transmembrane domain as they demonstrated the greatest change in structural orientation over the full GROMACS molecular dynamics (MD) simulation trajectory. The central α2-α4 regions of SPCA2C on the other hand adopt a stable packing, suggesting these domains may not require contacts with other proteins for stability. Importantly, deletion of the coiled-coiled domain of CCDC47 reduced or eliminated interactions with other SOCE proteins (i.e. STIM1 and Orai1, respectively) but did not alter interactions with SPCA2C. Remarkably however, further elimination of the entire cytosolic domain of CCDC47 restored interactions and CCDC47-STIM1 and -Orai1 interactions were maintained even after the deletion of both luminal and cytosolic domains of CCDC47. Elimination of the CCDC47 coiled-coil also perturbed ER localization of the protein. Only SPCA2C regularly localized to the plaques formed by DsRed2-CCDC47 ^MYC^, which supports the consistent interactions observed between the CCDC47 truncation and SPCA2C, but not with Orai1 or STIM1. The conserved coiled-coil domain in the CCDC47 cytosolic region is likely to participate in interactions that regulate intramolecular accessibility of the CCDC47 transmembrane domain and intermolecular interactions with the binding partners localized at the ER such as STIM1. It is also possible that the coiled-coil region functions to stabilize the cytosolic globular domain of the protein. Indeed, the coiled-coil of CCDC47 has an important role in SOCE regulation as re-expressing CCDC47 without the cytosolic domain does not restore SOCE to the same levels as the complete protein in *Ccdc47^−/−^* MEFs (24). Surprisingly, the ER transmembrane domain of CCDC47 was sufficient for interactions with Orai1, a plasma membrane protein. Therefore, CCDC47-Orai1 interactions appear to be bridged by other proteins located at ER-PM junctions.

It is possible that CCDC47 interacts with SPCA2C as part of its role as a chaperone for multi-spanning membrane proteins (35, 36). Given SPCA2C contains four transmembrane domains, CCDC47 which is part of the PAT complex, may play a role in the folding of SPCA2C (35). It is also possible that Orai1, a four transmembrane-spanning protein, is chaperoned by CCDC47. Conversely, SPCA2C may itself be involved with the PAT complex to assist in protein folding, a function critical to acinar cells. Again, future experiments will need to examine the PAT complex in situations where SPCA2C, CCDC47, or both are depleted.

In conclusion, this study defines the extensive SPCA2C interactome and reveals both known and unknown cellular roles. We reveal a network of SPCA2C-interacting proteins involved in mediating SOCE and provide insight into the molecular determinants of CCDC47 interactions with key SOCE regulators STIM1 and Orai1, as well as the pancreas-specific SPCA2C protein.

## Experimental Procedures

### Homology model of human SPCA2C

The Ca^2+^ ATPase structure in the RCSB Protein Data Bank showing the highest sequence similarity to human SPCA2C was used as a template to model the atomic-resolution, three-dimensional (3D) structure of SPCA2C. We modeled human SPCA2C because of a higher sequence similarity with any available experimentally determined template structure compared to mouse SPCA2C. MODELLER (version 9.16) was used to build the 3D models of human SPCA2C (37). We generated 20 homology models of human SPCA2C using 6LLE.pdb as the template structure (38). The model with the lowest discrete optimized protein energy (DOPE) score was taken as the highest quality model and used in subsequent MD simulations.

### Molecular dynamics (MD) simulations of SPCA2C in GROMACS

GROMACS Version 2020.4 was used to simulate changes in the 3D structure, folding and dynamics of our human SPCA2C model over 1 µs, with a time step of 2 fs (39). The protein was surrounded by a phospholipid bilayer to mimic the cell membrane. The CHARMM-GUI was used to build the lipid bilayer around human SPCA2C (40–42). The bilayer included 85 1-palmitoyl-2-oleoyl-sn-glycero-3-phosphocholine (POPC) lipids and 9 cholesterol molecules on one leaflet, and 84 POPC lipids and 8 cholesterol molecules on the second leaflet. 15,424 water molecules were added to the system to produce a 30 Å thick water layer on each side of the bilayer. POPC was chosen because this lipid is the main constituent of the ER membrane, and cholesterol was included to increase the fluidity of the bilayer (43). The system had an X:Y dimension ratio of 1 and was set to physiological temperature (310.15 K). The CHARMM36m forcefield was used to create the system. The system was electro-neutral without the addition of any counter ions. After number of particles, pressure and temperature (NPT) energy minimization and six-step equilibration (i.e. three steps of 125 ps and three steps of 500 ps), the production MD simulation was run for 1000 ns, and the trajectory was output with 10 ps resolution to assess structural and dynamical changes over time. Helix-helix distances, radius of gyration (Rg), root mean square fluctuation (RMSF) and order parameters (S2) were calculated using built-in GROMACS commands. PyMOL (version 2.5; Schrodinger, LLC) and VMD (version 1.9.4) were used to visualize structures and trajectories, respectively. All images were rendered using PyMOL (version 2.5).

### Cloning

An *MCS-SPCA2C-BirA*HA* expression vector was generated by ligating the *Atp2c2c* coding sequence into *Nhe1* and *BamHI* sites of *MCS-BirA*-HA* (Addgene, 74224). Mouse *Atp2c2c* was amplified using the *pcDNA3.1-SPCA2C-FLAG* expression vector as a template and *Atp2c2c* primers (20). To generate an *EGFP-SPCA2C-FLAG* vector, *Bgl2* and *Mfe1* restriction enzymes were used to remove the *ESYT1* sequence from the *EGFP-E-Syt1* vector (Addgene, 66830) and ligate the *Atp2c2c-FLAG* sequence amplified from the *pcDNA3.1-SPCA2C-FLAG* vector. To generate CCDC47 constructs, PCR-mediated site-directed mutagenesis was performed with the *pDsRed2-ER* mammalian expression vector (Clontech). A *BsiW1* restriction site was introduced by substituting adenine for guanine at the 1347^th^ nucleotide (*pDsRed2-ER-BsiW1mut*) and PCR amplified CCDC47 regions K21-M483, K21-N399, K21-N160, and N131-N160 was inserted using the *pEXPqcxip-hCCDC47-FLAG* vector (Addgene, 159141) as a template and primers designed to include a C-terminus MYC-tag. *BsiW1* and *EcoR1* were used to subclone the amplified CCDC47 truncations into separate *pDsRed2-ER-BsiW1mut* vectors. List of primers used for cloning and mutagenesis are in Supplementary Table S2. Plasmid sequences were verified by the London Regional Genomics Facility (Western University, London, ON).

### Cell Culture, transfection, and biotinylation

Human Embryonic Kidney (HEK) 293A cells or HEK293T cells with constitutive Orai1^YFP^ expression (HEK-Orai1^YFP^) were cultured at 37°C, 5% CO_2_ in Dulbecco’s Modified Eagle Medium (DMEM) containing high glucose supplemented with 10% FBS and 1% Pen-Strep (ThermoFisher). Culturing media for HEK-Orai1^YFP^ cells was additionally supplemented with 4 mg/ml G418 (ThermoFisher). For biotinylation experiments, HEK-Orai1^YFP^ cells were transfected at 70-80% confluence with *MCS-SPCA2C-BirA*-HA* or *MCS-BirA*-HA* using JetPrime Transfection Kit (Polyplus). 24 hours after transfection, cells were incubated for 24 hours with DMEM supplemented with 50 µM biotin. For co-IP and fluorescence imaging of recombinant proteins, the same protocol was performed as above to transfect *pEXPqcxip-hCCDC47-FLAG, pcDNA3.1-SPCA2C-mRFP, EGFP-SPCA2C-FLAG*, *pDsRed2-ER-BsiW1mut, pDsRed2-ER-BsiW1mut-CCDC47-K21-M483, -K21-N399, -K21-N160, -N131-N160, pcDNA3.1-SPCA2C-FLAG, pCMV6-XL5-STIM1* (Origene), or *pCMV6-STIM1-mCherry* (20, 44). Subsequent experiments were performed 24-30 hours post-transfection.

### Preparation of extracts for BioID

BioID assays were performed on three separate transfection experiments. For each replicate, four 75 cm^2^ flasks were prepared for transfection of either *MCS-SPCA2C-BirA*-HA* or control *MCS-BirA*-HA*. Twenty-four hours post-transfection, cells were washed in phosphate-buffered saline (PBS). Whole-cell extracts were prepared using lysis buffer (50 mM Tris-HCl pH 7.4, 500 mM NaCl, 0.2% SDS (w/v), 1 mg/mL leupeptin, 10 mg/mL aprotinin, 1 mg/mL pepstasin, 1 mM DTT). A final concentration of 2% (w/v) Triton X-100 was added and samples sonicated on ice twice for 30 seconds at output level 4 (Sonic Dismembrator Model 100, Ottawa, Ontario, Canada). 50 mM Tris-HCl pH 7.4 was added to double the volume, and samples subjected to another round of sonication. Samples were centrifuged for 10 minutes at 16,500 RCF at 4°C and supernatants transferred to new microcentrifuge tubes for streptavidin pull-down.

### Streptavidin pull-down of biotinylated proteins

Streptavidin-conjugated proteins were isolated as previously described (45). Briefly, whole cell extracts were incubated with equilibrated magnetic Dynabeads (MyOne Streptavidin C1; Invitrogen) overnight at 4°C. Beads were washed twice in wash buffer 1 (2% SDS w/v), once in wash buffer 2 (0.1% (w/v) deoxycholic acid, 1% (v/v) Triton X-100, 1 mM EDTA, 500 mM NaCl, 50 mM HEPES; pH 7.5) and once in wash buffer 3 (0.5% (w/v) deoxycholic acid, 0.5% (w/v) NP-40, 1 mM EDTA, 250 mM LiCl, 10 mM Tris-HCl; pH 7.4). A 50 µL aliquot was removed for western blot analysis, and remaining sample resuspended in 50 mM Tris-HCl, pH 7.4. Beads were re-centrifuged and resuspended in 50 mM ammonium bicarbonate.

### On-bead trypsin digests

Proteins bound to magnetic beads were reduced using 100 mM DTT for 30 minutes and alkylated at room temperature for 30 minutes in the dark with 1 M iodoacetamine (IAA; Sigma-Aldrich). Proteins were digested by agitating samples at 37°C for 4 hours with LysC (FUJIFILM Wako Pure Chemical Corporation, USA), overnight with Tryp-LysC (Promega), and then 4 hours in mass-spectrometry-grade trypsin (Promega). The missed cleavage rate for samples was an average of 3.9%-21.9%. Magnetic beads were centrifuged and pelleted with a magnet. Supernatants were dried by speed vacuum before resuspension in 0.1% trifluoroacetic acid (ThermoFisher). Samples were desalted using C18 Zip Tips (Sigma-Aldrich), dried by speed vacuum again and subsequently resuspended in 0.1% formic acid (ThermoFisher).

### Liquid chromatography (C) electrospray ionizing (ESI) tandem mass spectrometry (MS)

Samples were submitted to the University of Western Ontario Biological Mass Spectrometry Laboratory (BMSL, London, ON, Canada) for analysis by high-resolution LC-ESI-MS/MS. Peptides were identified using an ACQUITY M-Class UHPLC system (Waters) connected to an Orbitrap Elite mass spectrometer (ThermoFisher). Approximately 1 µg of samples was injected onto an ACUITY UPCL M-Class Symmetry C18 Trap Column, 5 µm x 2 mm, trapped for 6 minutes at a flow rate of 5 µL/ minute in 99% solution A (0.1% formic acid (FA) in water) and 1% solution B (0.1% FA in acetonitrile). ACUITY UPLC M-Class peptide BEH C18 columns (130 Å, 1.7 µm, 75 µm x 250 mm) were used to separate peptides. Separation was performed at a flow rate of 300 nL/minute at 7-23% solution B for 179 minutes, 23-35% solution B for 60 minutes, then 95% solution B and washing. Samples were run in positive ion mode.

### Mass spectrometry data analysis

Raw MS files were searched in PEAKS studio version X (Bioinformatics Solutions Inc., Waterloo, ON, Canada) using the Human Uniprot database to match peptides to protein sequences. Maximum missed cleavages were set to three fragment mass deviation at 20 ppm, fragment mass error tolerance was set to 0.8 Da. Protein and peptide false discover rate (FDR) was set to 1%. Cysteine carbamidomethylation was set as a fixed modification while oxidation (Met), amide deamidation (Asp/Gln), and acetylation (protein) were set as variable modifications with a maximum number of modifications per peptide set to 10. All other settings were left as default.

### Protein isolation and co-Immunoprecipitation

For protein isolation of membrane and cytosolic fractions, cells were harvested by trypsinization, rinsed with ice-cold PBS, and incubated first in a mild lysis buffer (150 mM NaCl, 50 nM HEPES pH 7.4, 25 µg/ml digitonin) for 10 minutes. Samples were centrifuged at 2,000 RCF and supernatant collected for the cytosolic protein fraction. The remaining insoluble pellet was washed with PBS, resuspended in a second lysis buffer (150 mM NaCl, 50 nM HEPES pH 7.4, 1% NP40), and incubated for 40 minutes with mild agitation. Samples were centrifuged at 7,000 RCF and the remaining supernatant was considered a membrane-enriched protein fraction. To isolate whole protein, cells were washed with ice-cold PBS and incubated at 4°C for 30 minutes with TNTE lysis buffer (30 mM Tris-HCl, 120 mM NaCl, 0.5% Triton X-100, 2.5 mM EDTA; pH 7.4). Samples were centrifuged at 21130 RCF for 10 minutes at 4°C. The supernatant was collected and used for immunoblotting, immunoprecipitation, and streptavidin pull-down experiments.

For co-IP experiments, 500 µg of whole protein lysates, membrane, or cytosolic fractions were incubated with 2 µL of primary antibody and 100 µL of Dynabeads™ Protein G (Invitrogen) overnight at 4°C. Proteins were immunoprecipitated with either mouse anti-HA (abcam), rabbit anti-Orai1 (Sigma), rabbit anti-ASPH (ThermoFisher), mouse anti-STIM1 (Novus), mouse anti-FLAG (Sigma), rabbit anti-CCDC47 (Sigma), or mouse anti-MYC (Cell Signaling). For experiments examining streptavidin pull-down of biotinylated ASPH, ESYT1, Orai1 and STIM1, 500 µg of whole protein was incubated with 100 µL superparamagnetic beads covalently coupled with streptavidin (MyOne Streptavidin C1; Invitrogen) overnight at 4°C.

### Immunoblotting

Cell lysates and immunoprecipitated protein were quantified using the Bradford assay (BioRad), resolved on 12% SDS-PAGE gels via electrophoresis, and transferred to a polyvinylidene difluoride (PVDF) membrane (BioRad). The membrane was incubated in blocking solution (5% non-fat skim milk in tris-buffered saline and 0.1% Tween 20 (TBST)) for 1 hour followed by incubation with primary antibody diluted in blocking solution overnight at 4°C. Mouse anti-HA (abcam), rabbit anti-Orai1 (Sigma), rabbit anti-ASPH (ThermoFisher), rabbit anti-ESYT1 (Sigma), mouse anti-STIM1 (Novus), mouse anti-FLAG (Sigma), rabbit anti-CCDC47 (Sigma) or rabbit anti-MYC (abcam) were used for WB analysis. Following several washes, membranes were incubated for 1 hour at room temperature with secondary anti-mouse or -rabbit-IgG HRP conjugated antibody (Cell Signalling) diluted in TBST. For visualization of biotinylated proteins, PVDF membrane with transferred protein was agitated in BSA blocking buffer (1% BSA (w/v) in 0.2% Triton X-100 (v/v) in PBS) for 20-30 minutes at room temperature, followed by incubation with HRP-conjugated streptavidin (abcam) diluted 1:10 000 in BSA blocking buffer for 40 minutes. Membrane was incubated for 5 minutes and then washed several times with BSA blocking buffer, prior to agitation in PBS for 5 minutes. BioRad Clarity Western ECL substrate was added to membranes to visualize protein using Bio-Rad VersaDoc Imaging System (BioRad) or iBright Imaging System (Thermofisher).

### Fluorescence Imaging

Transfected HEK293A or HEK-Orai1^YFP^ cells were fixed on coverslips in 4% (v/v) formalin or paraformaldehyde for 10 minutes at room temperature and then permeabilized with 0.1% Triton X-100 in PBS for 10-20 minutes. In preparation for immunofluorescent (IF) staining, cells were incubated for 30 minutes in blocking solution (5% BSA in 0.1% Triton X-100 (v/v) in PBS) and then for 1 hour at room temperature with respective primary antibody diluted in blocking solution. Mouse anti-STIM1 (Novus) or mouse anti-FLAG (Sigma) were used for IF staining. Cells were washed with PBS, and subsequently incubated for 1 hour with FITC-labeled secondary antibody (Jackson ImmunoResearch Laboratories) diluted in PBS. Biotinylated proteins were detected by incubating samples in Alexa Fluor 568 conjugated streptavidin (1:1000; ThermoFisher) diluted in blocking solution for 1 hour. Nuclei were visualized by adding DAPI (1:1000 in PBS; ThermoFisher) for 5 minutes and PermaFluor^TM^ (FisherScientific) was used to mount coverslips containing cells onto microscope slides. Epifluorescence microscopy images were acquired using Leica DM5500B Microscope (Leica microsystems, Concord, Ontario, Canada). Confocal images were acquired using Nikon Eclipse Ti2-E inverted microscope with X-Light V2 (CrestOptics) plug-in spinning disk and ORCA-Fusion Gen-III Digital sCMOS camera (C14440-20UP; Hamamatsu). A 60x/1.4NA oil objective was used to capture images.

### Image Analysis

To examine co-localization of SPCA2C^mRFP^ with CCDC47^FLAG^, line scan analysis was performed using NIS-Elements Advanced Research software (version 5.21.03). Vertical and perpendicular horizontal lines were drawn across the centre of each cell (defined as centre of the nucleus) and intensities along each line measured. Intensities were normalized to the maximum difference. Image J software (version 1.53t) (46) was used to subtract the average background value and the smooth function was applied to entire confocal images.

## Supporting information

This article contains supporting information.

## Abbreviations

Ca^2+^: Calcium
SOCE: store-operated calcium entry
ER: endoplasmic reticulum
STIM1: stromal interaction molecule 1
Orai1: calcium release-activated calcium modulator 1
SPCA2: secretory pathway calcium-ATPase 2
SICE: store-independent calcium entry
BioID: biotin labeling identification
CCDC47: coiled-coil domain-containing protein 47
SAINT: Significance Analysis of INTeractome
GO: Gene ontology
KEGG: Kyoto encyclopedia of genes and genomes
ESYT1: extended synaptotagmin-1
ASPH: aspartate beta-hydroxylase
HEK: human embryonic kidney
co-IP: co-immunoprecipitation
IF: immunofluorescence
SA: streptavidin affinity
MS: mass spectrometry

## Acknowledgments

This work was supported in part by grants provided by the Natural Sciences and Engineering Research Council and Children’s Health Foundation.

## References

1. Cremer, T., Neefjes, J., and Berlin, I. (2020) The journey of Ca2+ through the cell – Pulsing through the network of ER membrane contact sites. J. Cell Sci. 10.1242/jcs.249136

2. Dai, S., Venturini, E., Yadav, S., Lin, X., Clapp, D., Steckiewicz, M., Gocher-Demske, A. M., Grahame, D., and Edelman, A. M. (2022) Calcium/calmodulin-dependent protein kinase kinase 2 mediates pleiotropic effects of epidermal growth factor in cancer cells. BBA-Molecular Cell Res. 10.1016/j.bbamcr.2022.119252

3. Skrzypski, M., Kołodziejski, P. A., Mergler, S., Khajavi, N., Nowak, K. W., and Strowski, M. Z. (2016) TRPV6 modulates proliferation of human pancreatic neuroendocrine BON-1 tumour cells. Biosci. Rep. 36, 372

4. Thiel, G., Schmidt, T., and Rössler, O. G. (2021) Ca2+ microdomains, calcineurin and the regulation of gene transcription. Cells. 10.3390/cells10040875

5. Murphy, J. A., Criddle, D. N., Sherwood, M., Chvanov, M., Mukherjee, R., McLaughlin, E., Booth, D., Gerasimenko, J. V, Raraty, M. G. T., Ghaneh, P., Neoptolemos, J. P., Gerasimenko, O. V, Tepikin, A. V, Green, G. M., Reeve, J. R., Petersen, O. H., and Sutton, R. (2008) Direct Activation of Cytosolic Ca2+ Signaling and Enzyme Secretion by Cholecystokinin in Human Pancreatic Acinar Cells. Gastroenterology. 135, 632–641

6. Messenger, S. W., Falkowski, M. A., and Groblewski, G. E. (2014) Ca2+-regulated secretory granule exocytosis in pancreatic and parotid acinar cells. Cell Calcium. 55, 369–375

7. Wen, L., Voronina, S., Javed, M. A., Awais, M., Szatmary, P., Latawiec, D., Chvanov, M., Collier, D., Huang, W., Barrett, J., Begg, M., Stauderman, K., Roos, J., Grigoryev, S., Ramos, S., Rogers, E., Whitten, J., Velicelebi, G., Dunn, M., Tepikin, A. V., Criddle, D. N., and Sutton, R. (2015) Inhibitors of ORAI1 prevent cytosolic calcium-associated injury of human pancreatic acinar cells and acute pancreatitis in 3 mouse models. Gastroenterology. 149, 481–492.e7

8. Putney, J. W. (1986) A model for receptor-regulated calcium entry. Cell Calcium. 7, 1–12

9. Nieto-Felipe, J., Macias-Diaz, A., Sanchez-Collado, J., Berna-Erro, A., Jardin, I., Salido, G. M., Lopez, J. J., and Rosado, J. A. (2023) Role of Orai-family channels in the activation and regulation of transcriptional activity. J. Cell. Physiol. 238, 714–726

10. Navarro-Borelly, L., Somasundaram, A., Yamashita, M., Ren, D., Miller, R. J., and Prakriya, M. (2008) STIM1-Orai1 interactions and Orai1 conformational changes revealed by live-cell FRET microscopy. J. Physiol. 586, 5383–5401

11. Muik, M., Frischauf, I., Derler, I., Fahrner, M., Bergsmann, J., Eder, P., Schindl, R., Hesch, C., Polzinger, B., Fritsch, R., Kahr, H., Madl, J., Gruber, H., Groschner, K., and Romanin, C. (2008) Dynamic Coupling of the Putative Coiled-coil Domain of ORAI1 with STIM1 Mediates ORAI1 Channel Activation * □ S. 10.1074/jbc.M708898200

12. Liou, J., Fivaz, M., Inoue, T., and Meyer, T. (2007) Live-cell imaging reveals sequential oligomerization and local plasma membrane targeting of stromal interaction molecule 1 after Ca2+ store depletion. Proc. Natl. Acad. Sci. U. S. A. 104, 9301–9306

13. Albarran, L., Lopez, J. J., Amor, N. Ben, Martin-Cano, F. E., Berna-Erro, A., Smani, T., Salido, G. M., and Rosado, J. A. (2016) Dynamic interaction of SARAF with STIM1 and Orai1 to modulate store-operated calcium entry. Sci. Reports 2016 61. 6, 1–11

14. Selli, C., Erac, Y., Kosova, B., Erdal, E. S., and Tosun, M. (2015) Silencing of TRPC1 regulates store-operated calcium entry and proliferation in Huh7 hepatocellular carcinoma cells. Biomed. Pharmacother. 71, 194–200

15. Feng, M., Grice, D. M., Faddy, H. M., Nguyen, N., Leitch, S., Wang, Y., Muend, S., Kenny, P. A., Sukumar, S., Roberts-Thomson, S. J., Monteith, G. R., and Rao, R. (2010) Store-Independent Activation of Orai1 by SPCA2 in Mammary Tumors. Cell. 143, 84–98

16. Chantôme, A., Potier-Cartereau, M., Clarysse, L., Fromont, G., Marionneau-Lambot, S., Guéguinou, M., Pagés, J. C., Collin, C., Oullier, T., Girault, A., Arbion, F., Haelters, J. P., Jaffrés, P. A., Pinault, M., Besson, P., Joulin, V., Bougnoux, P., and Vandier, C. (2013) Pivotal role of the lipid raft SK3-orai1 complex in human cancer cell migration and bone metastases. Cancer Res. 73, 4852–4861

17. Peretti, M., Badaoui, M., Girault, A., Van Gulick, L., Mabille, M. P., Tebbakha, R., Sevestre, H., Morjani, H., and Ouadid-Ahidouch, H. (2019) Original association of ion transporters mediates the ECM-induced breast cancer cell survival: Kv10.1-Orai1-SPCA2 partnership. Sci. Rep. 10.1038/s41598-018-37602-7

18. Garside, V. C., Kowalik, A. S., Johnson, C. L., DiRenzo, D., Konieczny, S. F., and Pin, C. L. (2010) MIST1 regulates the pancreatic acinar cell expression of Atp2c2, the gene encoding secretory pathway calcium ATPase 2. Exp. Cell Res. 316, 2859–2870

19. Fenech, M. A., Sullivan, C. M., Ferreira, L. T., Mehmood, R., MacDonald, W. A., Stathopulos, P. B., and Pin, C. L. (2016) Atp2c2 Is Transcribed From a Unique Transcriptional Start Site in Mouse Pancreatic Acinar Cells. J. Cell. Physiol. 231, 2768– 2778

20. Fenech, M. A., Carter, M. M., Stathopulos, P. B., and Pin, C. L. (2020) The pancreas-specific form of secretory pathway calcium ATPase 2 regulates multiple pathways involved in calcium homeostasis. Biochim. Biophys. Acta - Mol. Cell Res. 1867, 118567

21. Xiang, M., Mohamalawari, D., and Rao, R. (2005) A novel isoform of the secretory pathway Ca2+,Mn 2+-ATPase, hSPCA2, has unusual properties and is expressed in the brain. J. Biol. Chem. 280, 11608–11614

22. Kim, D. I., Birendra, K. C., Zhu, W., Motamedchaboki, K., Doye, V., and Roux, K. J. (2014) Probing nuclear pore complex architecture with proximity-dependent biotinylation. Proc. Natl. Acad. Sci. U. S. A. 111, 2453–2461

23. Srikanth, S., Jew, M., Kim, K. Do, Yee, M. K., Abramson, J., and Gwack, Y. (2012) Junctate is a Ca2+-sensing structural component of Orai1 and stromal interaction molecule 1 (STIM1). Proc. Natl. Acad. Sci. U. S. A. 109, 8682–8687

24. Zhang, M., Yamazaki, T., Yazawa, M., Treves, S., Nishi, M., Murai, M., Shibata, E., Zorzato, F., and Takeshima, H. (2007) Calumin, a novel Ca 2+-binding transmembrane protein on the endoplasmic reticulum. Cell Calcium. 42, 83–90

25. Konno, M., Shirakawa, H., Miyake, T., Sakimoto, S., Nakagawa, T., and Kaneko, S. (2012) Calumin, a Ca2+-binding protein on the endoplasmic reticulum, alters the ion permeability of Ca2+ release-activated Ca2+ (CRAC) channels. Biochem. Biophys. Res. Commun. 417, 784–789

26. Thapa, K., Wu, K. C., Sarma, A., Grund, E. M., Szeto, A., Mendez, A. J., Gesta, S., Vishnudas, V. K., Narain, N. R., and Sarangarajan, R. (2018) Dysregulation of the calcium handling protein, CCDC47, is associated with diabetic cardiomyopathy. Cell Biosci. 8, 1–13

27. Kappel, S., Borgström, A., Stokłosa, P., Dörr, K., and Peinelt, C. (2019) Store-operated calcium entry in disease: Beyond STIM/Orai expression levels. Semin. Cell Dev. Biol. 94, 66–73

28. Sharma, A., Ramena, G. T., and Elble, R. C. (2021) Advances in intracellular calcium signaling reveal untapped targets for cancer therapy. Biomedicines. 10.3390/biomedicines9091077

29. Roux, K. J., Kim, D. I., Burke, B., and May, D. G. (2018) BioID: A Screen for Protein-Protein Interactions. Curr. Protoc. Protein Sci. 91, 19.23.1–19.23.15

30. Pin, C. L., Rukstalis, J. M., Johnson, C., and Konieczny, S. F. (2001) The bHLH transcription factor Mist1 is required to maintain exocrine pancreas cell organization and acinar cell identity. J. Cell Biol. 155, 519–530

31. Chang, C. L., Hsieh, T. S., Yang, T. T., Rothberg, K. G., Azizoglu, D. B., Volk, E., Liao, J. C., and Liou, J. (2013) Feedback regulation of receptor-induced ca2+ signaling mediated by e-syt1 and nir2 at endoplasmic reticulum-plasma membrane junctions. Cell Rep. 5, 813–825

32. Kang, F., Zhou, M., Huang, X., Fan, J., Wei, L., Boulanger, J., Liu, Z., Salamero, J., Liu, Y., and Chen, L. (2019) E-syt1 Re-arranges STIM1 Clusters to Stabilize Ring-shaped ER-PM Contact Sites and Accelerate Ca 2+ Store Replenishment. Sci. Rep. 10.1038/s41598-019-40331-0

33. Dinchuk, J. E., Focht, R. J., Kelley, J. A., Henderson, N. L., Zolotarjova, N. I., Wynn, R., Neff, N. T., Link, J., Huber, R. M., Burn, T. C., Rupar, M. J., Cunningham, M. R., Selling, B. H., Ma, J., Stern, A. A., Hollis, G. F., Stein, R. B., Friedman, P. A., Dinchuk, J. E., Focht, R. J., Kelley, J. A., Henderson, N. L., Huber, R. M., Burn, T. C., Hollis, G. F., Stein, R. B., and Friedman, P. A. (2002) Absence of Post-translational Aspartyl-Hydroxylation of Epidermal Growth Factor Domains in Mice Leads to Developmental Defects and an Increased Incidence of Intestinal Neoplasia*. J. Biol. Chem. 277, 12970–12977

34. Morimoto, M., Waller-Evans, H., Ammous, Z., Song, X., Strauss, K. A., Pehlivan, D., Gonzaga-Jauregui, C., Puffenberger, E. G., Holst, C. R., Karaca, E., Brigatti, K. W., Maguire, E., Coban-Akdemir, Z. H., Amagata, A., Lau, C. C., Chepa-Lotrea, X., Macnamara, E., Tos, T., Isikay, S., Nehrebecky, M., Overton, J. D., Klein, M., Markello, T. C., Posey, J. E., Adams, D. R., Lloyd-Evans, E., Lupski, J. R., Gahl, W. A., and Malicdan, M. C. V. (2018) Bi-allelic CCDC47 Variants Cause a Disorder Characterized by Woolly Hair, Liver Dysfunction, Dysmorphic Features, and Global Developmental Delay. Am. J. Hum. Genet. 103, 794–807

35. Chitwood, P. J., and Hegde, R. S. (2020) An intramembrane chaperone complex facilitates membrane protein biogenesis. Nature. 584, 630–634

36. McGilvray, P. T., Anghel, S. A., Sundaram, A., Zhong, F., Trnka, M. J., Fuller, J. R., Hu, H., Burlingame, A. L., and Keenan, R. J. (2020) An ER translocon for multi-pass membrane protein biogenesis. Elife. 9, 1–43

37. Webb, B., and Sali, A. (2017) Comparative Protein Structure Modeling Using MODELLER. Curr. Protoc. Bioinforma. 1654, 39–54

38. Zhang, Y., Inoue, M., Tsutsumi, A., Watanabe, S., Nishizawa, T., Nagata, K., Kikkawa, M., and Inaba, K. (2020) Cryo-EM structures of SERCA2b reveal the mechanism of regulation by the luminal extension tail. Sci. Adv. 6, 147–159

39. Abraham, M. J., Murtola, T., Schulz, R., Páll, S., Smith, J. C., Hess, B., and Lindahl, E. (2015) ScienceDirect GROMACS: High performance molecular simulations through multi-level parallelism from laptops to supercomputers. SoftwareX. 1-2 (C), 19–25

40. Jo, S., Kim, T., and Im, W. (2007) Automated Builder and Database of Protein/Membrane Complexes for Molecular Dynamics Simulations. PLoS One. 2, e880

41. Jo, S., Kim, T., Iyer, V. G., and Im, W. (2008) CHARMM-GUI: A web-based graphical user interface for CHARMM. J. Comput. Chem. 29, 1859–1865

42. Lee, J., Cheng, X., Swails, J. M., Yeom, M. S., Eastman, P. K., Lemkul, J. A., Wei, S., Buckner, J., Jeong, J. C., Qi, Y., Jo, S., Pande, V. S., Case, D. A., Brooks, C. L., Mackerell, A. D., Klauda, J. B., and Im, W. (2016) CHARMM-GUI Input Generator for NAMD, GROMACS, AMBER, OpenMM, and CHARMM/OpenMM Simulations Using the CHARMM36 Additive Force Field. J. Chem. Theory Comput. 12, 405–413

43. Van Meer, G., and De Kroon, A. I. P. M. (2011) Lipid map of the mammalian cell. J. Cell Sci. 124, 5–8

44. Luik, R. M., Wu, M. M., Buchanan, J. A., and Lewis, R. S. (2006) The elementary unit of store-operated Ca2+ entry: Local activation of CRAC channels by STIM1 at ER-plasma membrane junctions. J. Cell Biol. 174, 815–825

45. Mehus, A. A., Anderson, R. H., and Roux, K. J. (2016) BioID Identification of Lamin-Associated Proteins. in Methods in Enzymology, pp. 3–22, 569, 3–22

46. Schneider, C. A., Rasband, W. S., and Eliceiri, K. W. (2012) NIH Image to ImageJ: 25 years of image analysis. Nat. Methods. 10.1038/nmeth.2089

47. Yao, X., and Xue, Y. (2009) LETTER TO THE EDITOR DOG 1.0: illustrator of protein domain structures. Mol. Biol. Cell Mol. Cel-lular Biol. 19, 271–273

