## Supplemental Figures for "Defining the SPCA2C interactome identifies unique links to Store-Operated Ca^2+^ Entry"

#### Figure Legends

**Figure S1. SPCA2C localization is not altered by BirA* enzyme fusion and is site of biotinylation within the cell.** (**A**) Immunofluorescence for fluorescein-stained HA (green) was combined with fluorescence of mRFP following transient co-expression of SPCA2C-BirA$^{*HA}$ and SPCA2C$^{mRFP}$ in HEK293 cells. Arrows indicate areas of overlap. (**B**) Combined fluorescence for IF-stained HA (red) and Orai1$^{YFP}$ (green) in HEK-Orai1$^{YFP}$ cells with transient expression of BirA$^{*HA}$ (negative control; top panel) or SPCA2C-BirA$^{*HA}$ (bottom panel) . Arrows indicate areas of Orai1$^{YFP}$ localization. The area of overlap in fluorescence signal is greater between Orai1$^{YFP}$ and SPCA2C-BirA$^{*HA}$ (bottom panel) compared to BirA$^{*HA}$ (top panel). (**C**) IF for HA (green) was combined with visualization of biotinylated proteins using Alexa Fluor 568- conjugated streptavidin (red) following transient expression of BirA$^{*HA}$ (top panel) or SPCA2C-BirA$^{*HA}$ (bottom panel) in HEK293 cells. Cells were incubated with 50 µM of biotin for 24 hours prior to fixation and visualization. Arrows indicate regions of overlap. Localization of biotinylated proteins is distinct in SPCA2C-BirA$^{*HA}$ expressing cells compared to cells with BirA$^{*HA}$ expression. Scale bars= 10 µm. Nuclei stained with DAPI (blue).

**Figure S2. Schematic showing analysis and stringency in defining the SPCA2C interactome.** (**A**) Biotinylated proteins from whole protein lysates of either BirA$^{*HA}$ (negative control) or SPCA2C-BirA$^{*HA}$ expressing Hek-Orai1$^{YFP}$ cells incubated 24 hours with biotin were collected via affinity capture and identified using MS analysis from three independent experiments. Proteins that appeared in both SPCA2C-BirA$^{*HA}$ and BirA$^{*HA}$ MS lists or those that did not appear in all three SPCA2C-BirA$^{*HA}$ MS lists were omitted, unless they received a SAINT score greater than or equal to 0.95, suggesting a true interactor. Number of proteins listed in each step is included in brackets. (**B**) Western blot analysis for biotinylated proteins detected via Streptavidin-HRP from BirA$^{*HA}$ or SPCA2C-BirA$^{*HA}$-expressing HEK-Orai1$^{YFP}$ cells.

**Figure S3. Cluster analysis on the SPCA2C interactome indicates ER and vesicular transport-related functions.** A protein-protein interaction network of candidate SPCA2C-interacting proteins from BioID assay was established by STRING and groups of proteins were clustered by the ClusterOne plug-in in Cytoscape. Groups were classified based on GO Biological Process, Molecular Function, or KEGG pathway analysis of collective proteins in each group.

**Figure S4. Single and merged channel confocal images of DsRed2- CCDC47 constructs with SPCA2C, STIM1, or Orai1.** Spinning disk confocal microscopy visualization of DsRed2 fused to calreticulin signal sequence (control; DsRed2-ER), CCDC47$_{full}$, CCDC47$_{\Delta CC}$, CCDC47$_{\Delta G+\Delta CC}$, or CCDC47$_{TM}$ (red) with (**A**) EGFP-SPCA2C$^{FLAG}$, (**B**) immunofluorescent labeled STIM1, or (**C**) Orai1$^{YFP}$ (green) in HEK293 cells. Areas of colocalization of CCDC47 constructs with SPCA2C, STIM1, or Orai1 are indicated with white arrows. Single-plane images at 0.3 µm thickness are shown. Scale bar=10 µm.

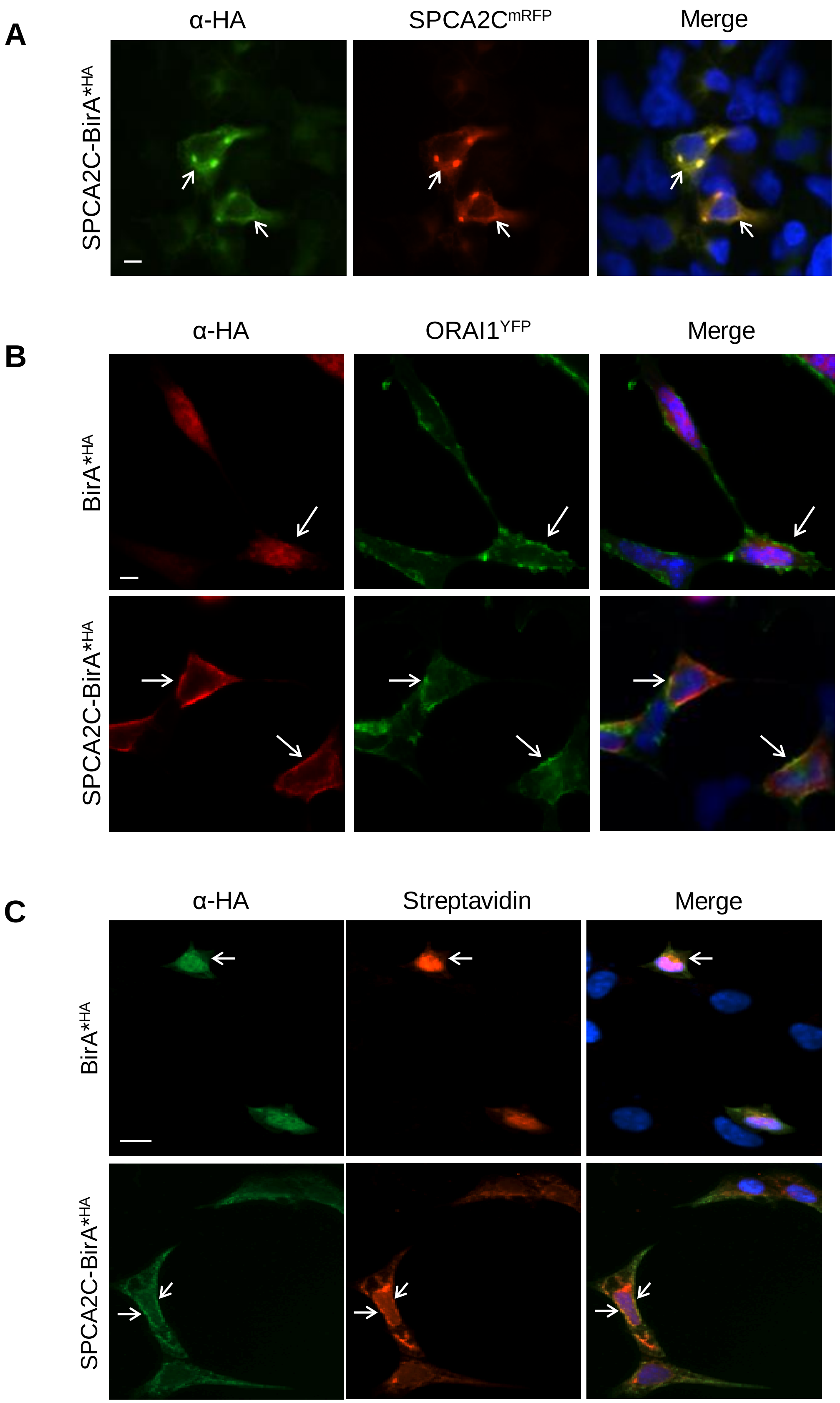


**Samardzija et al (2023) Figure S1**

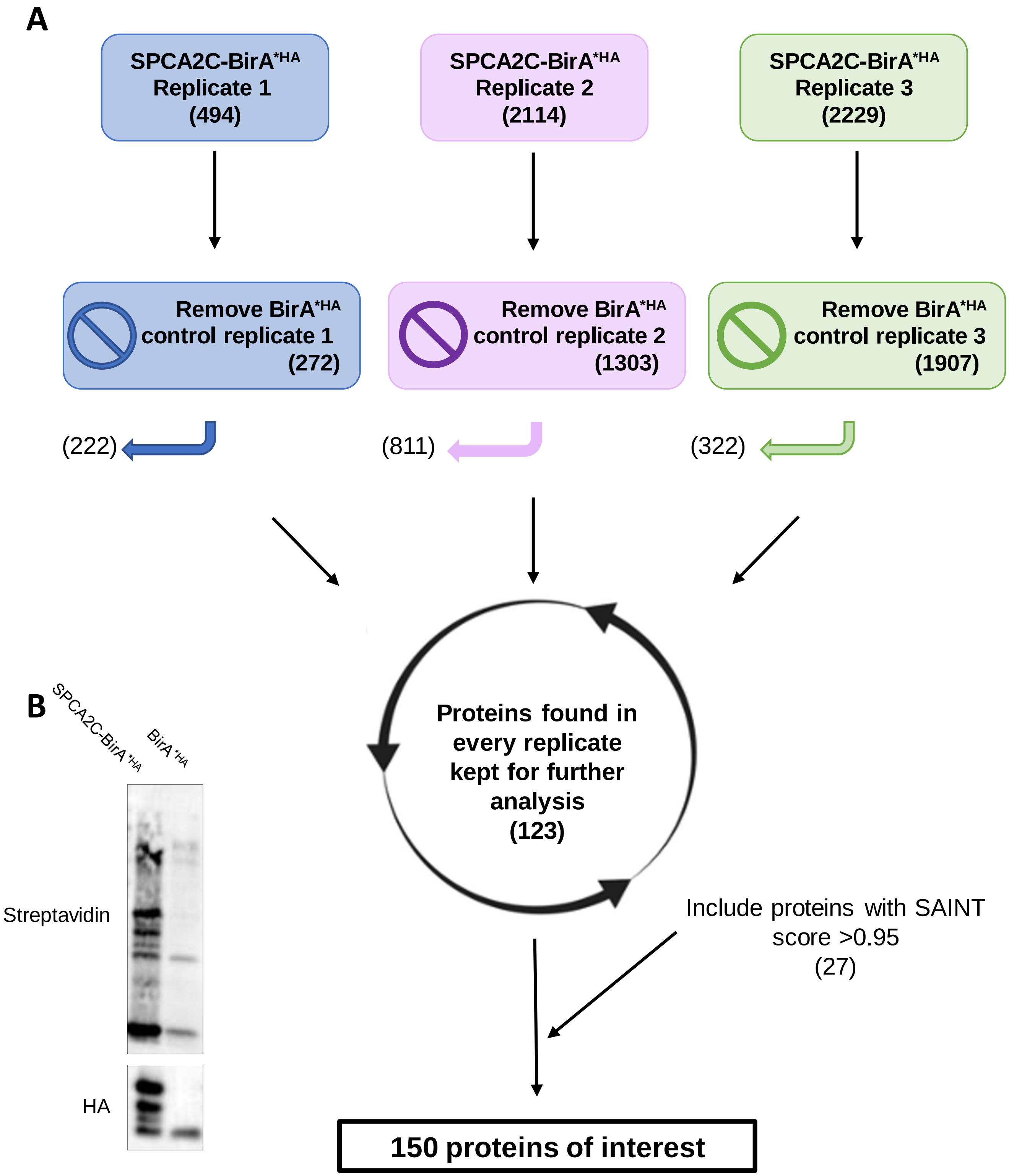


**Samardzija et al (2023) Figure S2**

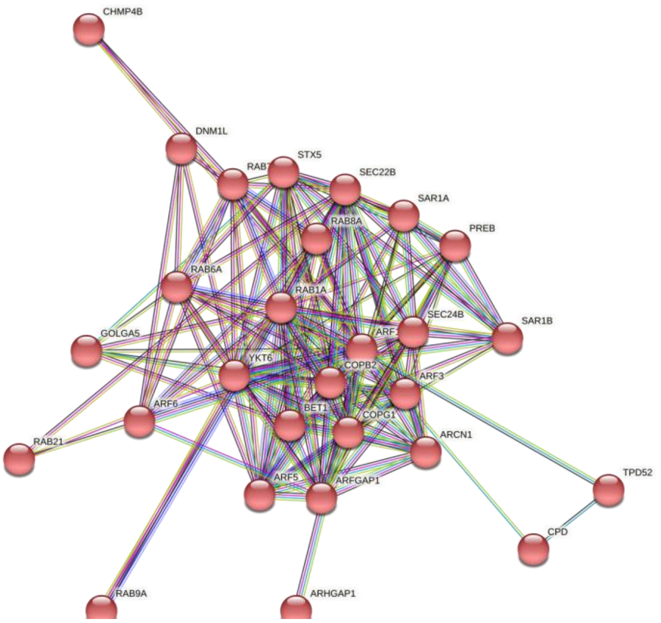


SNARE interactions in vesicular transport

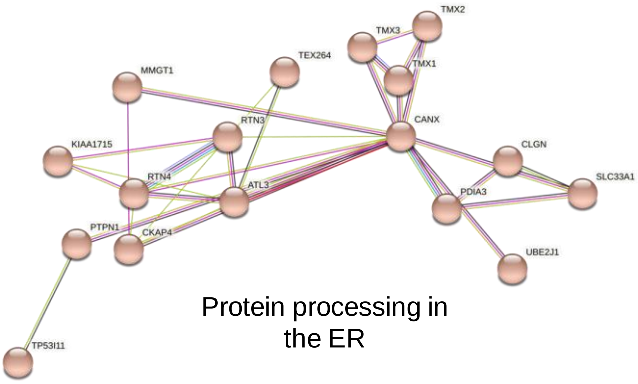


Protein processing in the ER

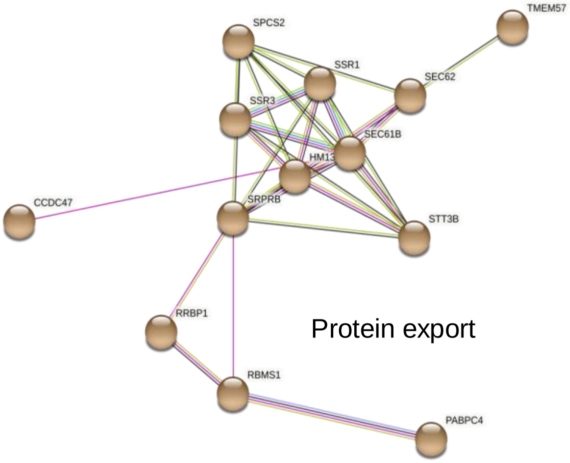


Protein export

**Samardzija et al (2023) Figure S3**

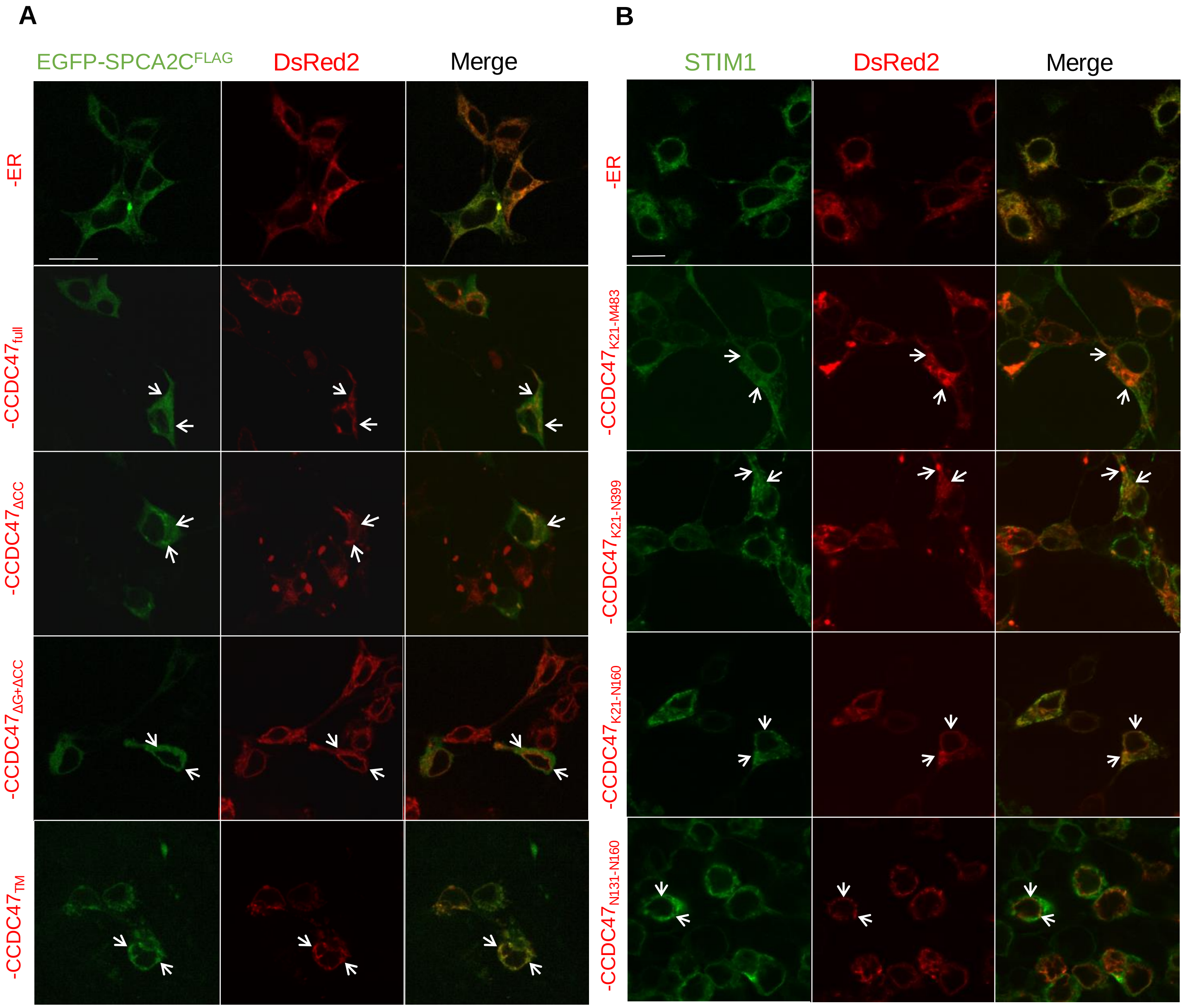


**Samardzija et al (2023) Figure S4**

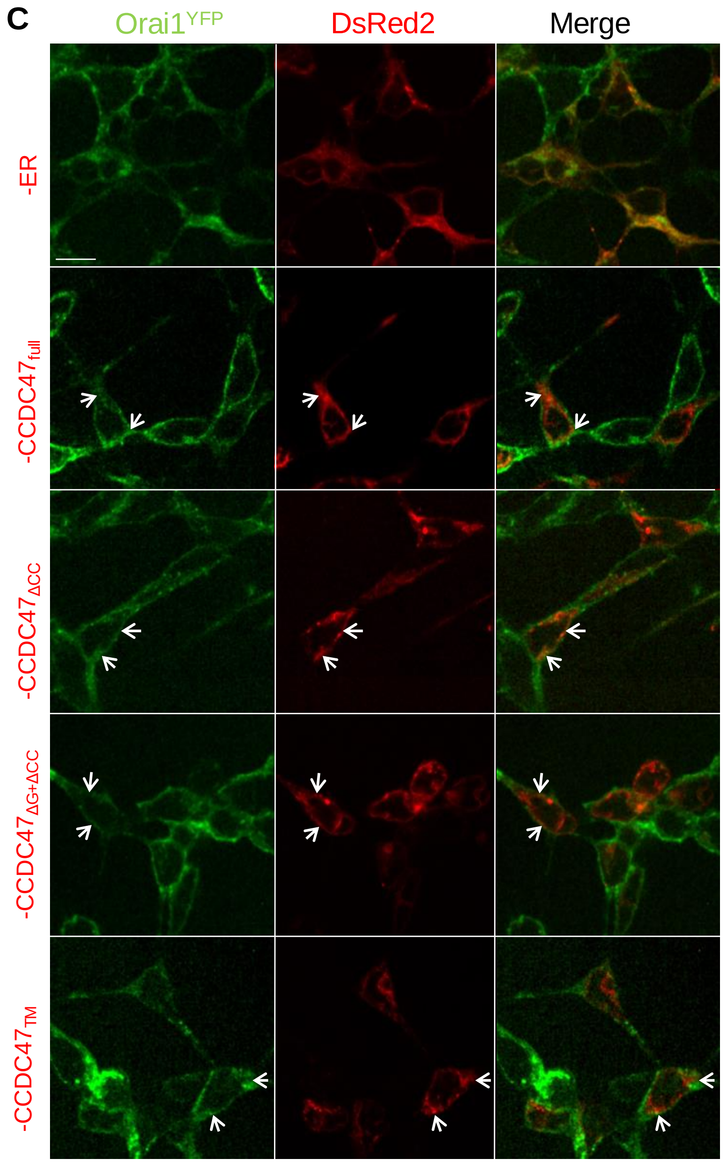


Samardzija et al (2023) Figure S4

### Table S1. SAINT analysis

| Protein ID | Gene | Max FCA | Related to $Ca^{2+}$ | Disease |
|---|---|---|---|---|
| CKAP4 | *CKAP4* | 26.32 | No | Cystitis and Axenfeld-Rieger Syndrome |
| CCD47 | *CCDC47* | 25.62 | Yes | Trichohepatoneurodevelopmental Syndrome; Atypical Choroid Plexus Papilloma |
| UBXN4 | *UBXN4* | 25.14 | No | Inclusion Body Myopathy;Martsolf Syndrome 1 |
| VAPB | *VAPB* | 19.78 | Yes | Amyotrophic Lateral Sclerosis;Spinal Muscular Atrophy, Late-Onset, Finkel Type |
| TMX1 | *TMX1* | 19.02 | No | |
| PTN1 | *PTPN1* | 18.7 | No | Type 2 Diabetes Mellitus |
| 4F2 | *SLC3A2* | 17.67 | No | Lysinuric Protein Intolerance; Cystinuria |
| ITB1 | *ITGB1* | 15.84 | Yes | Gallbladder and Breast Cancer |
| HM13 | *HM13* | 15.53 | No | Hepatitis C |
| STX5 | *STX5* | 15.41 | No | Human Cytomegalovirus Infection; Endometriosis |
| KTN1 | *KTN1* | 14.86 | No | Gliofibroma and Hepatocellular Carcinoma. |
| RRBP1 | *RRBP1* | 14.58 | No | Epithelioid Inflammatory Myofibroblastic Sarcoma |
| VAPA | *VAPA* | 12.99 | No | |
| TOIP1 | *TOR1A1IP1* | 11.83 | No | Myopathy; Muscular Dystrophy |
| DNM1L | *DNM1L* | 11.48 | No | Encephalopathy |
| LYRIC | *MTDH* | 11.34 | No | Glioblastoma and Hepatocellular Carcinoma |
| HXK1 | *HXK1* | 11.15 | No | Hemolytic Anemia |
| CALX | *CANX* | 10.92 | Yes | Influenza; Nephrogenic Diabetes Insipidus |
| STT3B | *STT3B* | 10.53 | No | Immunodeficiency, X-Linked, |
| ARF1 | *ARF1* | 9.99 | No | Periventricular Nodular Heterotopia |
| ARF3 | *ARF3* | 9.99 | No | Pectus Excavatum; Scoliosis |
| MAN1 | *LEMD3* | 9.73 | No | Buschke-Ollendorff Syndrome |
| STIM1 | *STIM1* | 9.63 | Yes | Stormorken Syndrome |
| RTN4 | *RTN4* | 9.27 | No | Temporal Lobe Epilepsy; Anosognosia. |
| ARF5 | *ARF5* | 9.27 | No | Cholera and Geroderma Osteodysplasticum. |
| HMOX2 | *HMOX2* | 9.09 | No | |
| RAB1A | *RAB1A* | 8.87 | No | Choroideremia and Spinocerebellar Ataxia |
| LRC59 | *LRRC59* | 8.57 | No | Ogden syndrome |
| NB5R1 | *CYB5R1* | 8.43 | Yes | De Quervain Disease |
| CLCC1 | *CLCC1* | 8.35 | No | Retinitis Pigmentosa 32 and Retinitis |
| ASPH | *ASPH* | 8.19 | Yes | Facial Dysmorphism, Lens Dislocation, |
| DHRS7 | *DHRS7* | 8.17 | No | |
| RAB8A | *RAB8A* | 8.11 | No | Microvillus Inclusion Disease; Glaucoma |
| PGRC1 | *PGRC1* | 7.9 | No | Extragonadal Germ Cell Cancer |
| PDIA3 | *PDIA3* | 7.76 | ? | Macular Degeneration; Gastric Cancer |
| ANKL2 | *ANKL2* | 7.67 | No | Microcephaly |
| LBR | *LBR* | 7.28 | No | Greenberg Dysplasia; Rhizomelic Skeletal Dysplasia |
| RHG01 | *ARHGAP1* | 7.18 | No | Nephrogenic Diabetes Insipidus; Lowe Oculocerebrorenal Syndrome |
| SAR1A | *SAR1A* | 7.13 | No | Spondyloepiphyseal Dysplasia Tarda; Chylomicron Retention Disease |
| RAB6A | *RAB6A* | 7.04 | No | Cohen Syndrome; Choroideremia |
| EMD | *EMD* | 6.99 | no | Emery-Dreifuss Muscular Dystrophy; X-Linked Emery-Dreifuss Muscular Dystrophy |
| CP51A | *CYP51A1* | 6.67 | no | Chagas Disease |
| TM109 | *TMEM109* | 6.54 | Yes | |
| F1142 | *F1142* | 6.4 | No | Isolated Ectopia Lentis |
| YKT6 | *YKT6* | 6.22 | No | |
| ESYT1 | *ESYT1* | 6.15 | Yes | Congenital Nystagmus 1; Stormorken Syndrome. |
| RAB7A | *RAB7A* | 6.12 | No | Charcot-Marie-Tooth Disease |
| HSPB1 | *HSPB1* | 6.09 | No | Charcot-Marie-Tooth Disease |
| SRPRB | *SRPRB* | 6.09 | No | Testis Seminoma |
| RAB9A | *RAB9A* | 6.08 | No | Warburg Micro Syndrome 1 |
| PHLP | *PDCL* | 6.06 | No | Pollen Allergy and Timothy Grass Allergy |
| IF4H | *EIF4H* | 6.01 | No | Williams-Beuren Syndrome |
| SSRA | *SSRA* | 5.78 | ? | |
| UB2J1 | *UBE2J1* | 5.66 | No | Motor Neuritis |

| Protein ID | Gene | Max FCA | Related to $Ca^{2+}$ | Disease |
|---|---|---|---|---|
| CLGN | *CLGN* | 5.51 | Yes | Colloid Adenoma |
| MAVS | *MAVS* | 5.46 | No | Mouth Disease; Hepatitis |
| PGRC2 | *PGRC2* | 5.44 | No | Schizophrenia; Achalasia-Addisonianism-Alacrima Syndrome |
| SQSTM | *SQSTM* | 5.11 | No | Paget Disease; Frontotemporal Dementia; Amyotrophic Lateral Sclerosis |
| SAR1B | *SAR1B* | 5 | No | Chylomicron Retention Disease; Hypobetalipoproteinemia, Familial |
| MXRA7 | *MXRA7* | 4.81 | No | |
| TX264 | *TX264* | 4.71 | Yes | |
| DHCR7 | *DHCR7* | 4.6 | No | Smith-Lemli-Opitz Syndrome |
| BET1 | *BET1* | 4.44 | No | Citrullinemia, Type II |
| SC22B | *SEC22B* | 4.32 | No | Legionellosis |
| HACD3 | *HACD3* | 4.21 | No | Spinocerebellar Ataxia; Fiedler's Myocarditis |
| ZYX | *ZYX* | 4.2 | No | Lipomatosis |
| IF5A1 | *EIF5A* | 4.09 | No | Faundes-Banka Syndrome |
| VAS1 | *VARS1* | 4.02 | No | Neurodevelopmental Disorder |
| RABL3 | *RABL3* | 3.99 | No | Pancreatic cancer |
| MBOA7 | *MBOAT7* | 3.79 | No | Intellectual Developmental Disorder, |
| NCPR | *POR* | 3.6 | No | Antley-Bixler Syndrome |
| PCBP1 | *PCBP1* | 3.34 | No | Deafness |
| PROF1 | *PFN1* | 3.28 | Yes | Amyotrophic Lateral Sclerosis |
| TPD54 | *TPD52L2* | 3.08 | No | Breast cancer |
| FHL1 | *FHL1* | 2.62 | No | Myopathy, X-Linked |
| DNJB1 | *DNAJB1* | 2.33 | No | Fibrolamellar Carcinoma; Muscular Dystrophy, |
| TERA | *SINHCAF* | 7.6 | No | |
| COPG1 | *COPG1* | 5.64 | No | Parainfluenza Virus; Listeria Meningitis. |
| SPCS2 | *SPCS2* | 5.63 | No | Epiphyseal Dysplasia |
| CDKAL | *CDKAL* | 5.47 | No | Type 2 Diabetes Mellitus |
| STBD1 | *STBD1* | 4.71 | No | Congenital and Glycogen Storage Disease II |
| LNP | *LNPK* | 4.71 | No | Neurodevelopmental Disorder with Epilepsy; Robinow Syndrome, Autosomal Recessive |
| ITM2B | *ITM2B* | 4.17 | No | Retinal Dystrophy; Cerebral Amyloid Angiopathy |
| MTX3 | *MTX3* | 4.06 | No | Loeys-Dietz Syndrome 1; Transient Arthritis |
| TFCP2 | *TFCP2* | 3.89 | No | Rhabdomyosarcoma |
| AT2A2 | *ATP2A2* | 3.78 | Yes | Darier-White Disease |
| RCN1 | *RCN1* | 3.76 | Yes | Wilms Tumor; Impaired Intellectual Development |
| MMGT1 | *MMGT1* | 3.74 | No | Fundus Dystrophy |
| S38A2 | *SLC38A2* | 3.64 | No | Hartnup Disorder |
| RFC4 | *RFC4* | 3.57 | No | Fanconi Anemia; Gastric Cancer |
| DHB12 | *HSD17B12* | 3.37 | No | Cervical Neuroblastoma |
| TMX2 | *TMX2* | 3.32 | No | Neurodevelopmental Disorder |
| AAAT | *SLC1A5* | 3.29 | No | Hartnup Disorder |
| 1433T | *YWHAQ* | 3.29 | No | Alzheimer Disease |
| ARFG1 | *ARFGAP1* | 3.04 | No | Ceroid Lipofuscinosis; Hypotrichosis-Lymphedema-Telangiectasia Syndrome |
| ALG9 | *ALG9* | 3.02 | No | Congenital Disorder of Glycosylation, Type II; Gillessen-Kaesbach-Nishimura Syndrome |
| TAGL2 | *TAGLN2* | 2.7 | Yes | Barrett's Adenocarcinoma; Maxillary Sinus Squamous Cell Carcinoma |
| SSRG | *SSR3* | 2.62 | Yes | Congenital Disorder of Glycosylation; Esophageal Cancer |
| EI24 | *EI24* | 2.56 | No | |
| DDX17 | *DDX17* | 6.67 | No | |
| LAP2B | *TMPO* | 4.93 | No | Dilated Cardiomyopathy |
| H2B3B | *H2BC26* | 4.58 | No | Spherocytosis, Type 1 |
| RBMS1 | *RBMS1* | 3.98 | No | Diffuse Glomerulonephritis and Microcephaly 2 |
| CHM4B | *CHMP4B* | 3.28 | No | Cataracts |
| F177A | *FAM177A1* | 2.3 | No | Myasthenic Syndrome; Hypotonia-Cystinuria Syndrome |
| STIP1 | *STIP1* | 4.08 | No | Microcephaly Recessive; Chronic Wasting Disease |
| COPD | *COPD* | 3.41 | No | Pulmonary Disease; Chronic Obstructive |
| S38A1 | *SLC38A1* | 2.25 | No | Klatskin's Tumor |

[1]Ranks given from 108 proteins based on FC score; [2]Fold change (FC): Based on the ratio of average normalized spectral counts in bait purifications to negative controls. Higher the number indicates a more likely interaction; [3]Significance Analysis of Interactome (SAINT): Provides a statistical conversion of the FC score onto the probability scale using the mixed-model analysis of underlying spectral count distribution; [4]Function based on literature review. Gray rows indicate those proteins identified through SAINT but not in all three BioID screens.

**Table S2. Primer sequences used for cloning**

| **Primer** | **Purpose** | **Sequence (5'-3')** |
|---|---|---|
| BsiW1-mutation | Site-directed mutagenesis of *pDsRed2-ER* | Forward: CCTGTTCCTGAGATCGTACGAGAAGGACGAGC<br>Reverse: GCTCGTCCTTCTCGTACGATCTCAGGAACAGG |
| *Atp2c2c* | Cloning into MCS-BirA*-HA | Forward: CTAGGCTAGCGACGATCCTGAACAGA<br>Reverse: AGTCGAGGATCCCTCCACGGAGTC |
| *Atp2c2c* | Cloning into *pEGFP-C1* | Forward: CCTGATCCTGAGGAGATCTATGTCGGCCGCTG<br>Reverse: GCTGGATATCTGCCAATTGCCACCACACTGGAC |
| $CCDC47_{K21}$-Forward | Cloning into *pDsRed2-ER-BsiW1mut* | GTGTCTCTCGTACGGTAAGTTTGATGATTTTGAGGATGAGG |
| $CCDC47_{N131}$-Forward | Cloning into *pDsRed2-ER-BsiW1mut* | CCTGCACACCGTACGGTAACAGCTGGGAGAG |
| $CCDC47_{M483}$-Reverse | Cloning into *pDsRed2-ER-BsiW1mut* | TCATCGAATTCTTACAGATCCTCTTCTGAGATGAGTTTTTGTT CCATGGCTTTCACTTTGATTTG |
| $CCDC47_{N399}$-Reverse | Cloning into *pDsRed2-ER-BsiW1mut* | TCTACGAATTCTCACAGATCCTCTTCTGAGATGAGTTTTTGT TCGTTGAGTCGGAACTTTTTGGC |
| $CCDC47_{N160}$-Reverse | Cloning into *pDsRed2-ER-BsiW1mut* | CAAAAGGAATTCTCACAGATCCTCTTCTGAGATGAGTTTTTG TTCGTTTTTATTCTTCCCAATG |
